# Adipocyte Leptin Signaling Regulates Glycemia and Cardiovascular Function via Enhancing Brown Adipose Tissue Thermogenesis in Obese Male Mice

**DOI:** 10.1101/2025.10.10.681380

**Authors:** Yoichi Ono, Simone Kennard, Benjamin T. Wall, Jing Ma, Eric J. Belin de Chantemèle

## Abstract

While leptin control of metabolism is primarily viewed as centrally mediated, leptin has also been shown to directly regulate adipocyte function. However, the impact of the peripheral effects of leptin on systemic metabolism, especially in the context of obesity, remains unclear. To address this question, we selectively restored adipocyte leptin receptor (LEPR) expression in obese male and female LEPR conditional KO mice. Adipocyte LEPR restoration did not affect body weight but selectively increased brown adipose tissue (BAT) mass in male mice. This was associated with increased energy expenditure, smaller BAT adipocytes, lower triglycerides content, and increased markers of browning and lipolysis exclusively in males. Additionally, adipocyte LEPR restoration enhanced the expression of markers of endothelial cell and angiogenesis in male BAT, supporting increased local vascularization. Improved BAT function in males was also associated with lower HbA1c, better insulin sensitivity, reduced systolic blood pressure, decreased arterial stiffness and improved endothelial function. Lastly, adipocyte LEPR restoration lowered circulating pro-inflammatory cytokines and reduced tissue inflammation in the aorta and the heart, again in males only. These findings reveal a critical role for adipocyte leptin signaling in regulating BAT function and emphasize its importance in maintaining glycemic and cardiovascular health in males with obesity.

**Article Highlights:** Leptin is known to enhance BAT activity through sympathetic stimulation. However, *in vitro* studies suggest leptin could also act directly on adipocytes to promote lipolysis. Whether these peripheral effects of leptin are relevant to systemic metabolic control, in obesity, remain ill-defined. We addressed this question by selectively restoring leptin receptor (LEPR) in adipocytes of obese LEPR conditional KO mice. LEPR restoration selectively enhanced BAT activity in male mice, which led to improved glycemic control and cardiovascular function. These findings revealed a crucial role for BAT leptin signaling in regulating energy expenditure, glycemic and cardiovascular health, primarily in males.

**Graphical Abstract:** 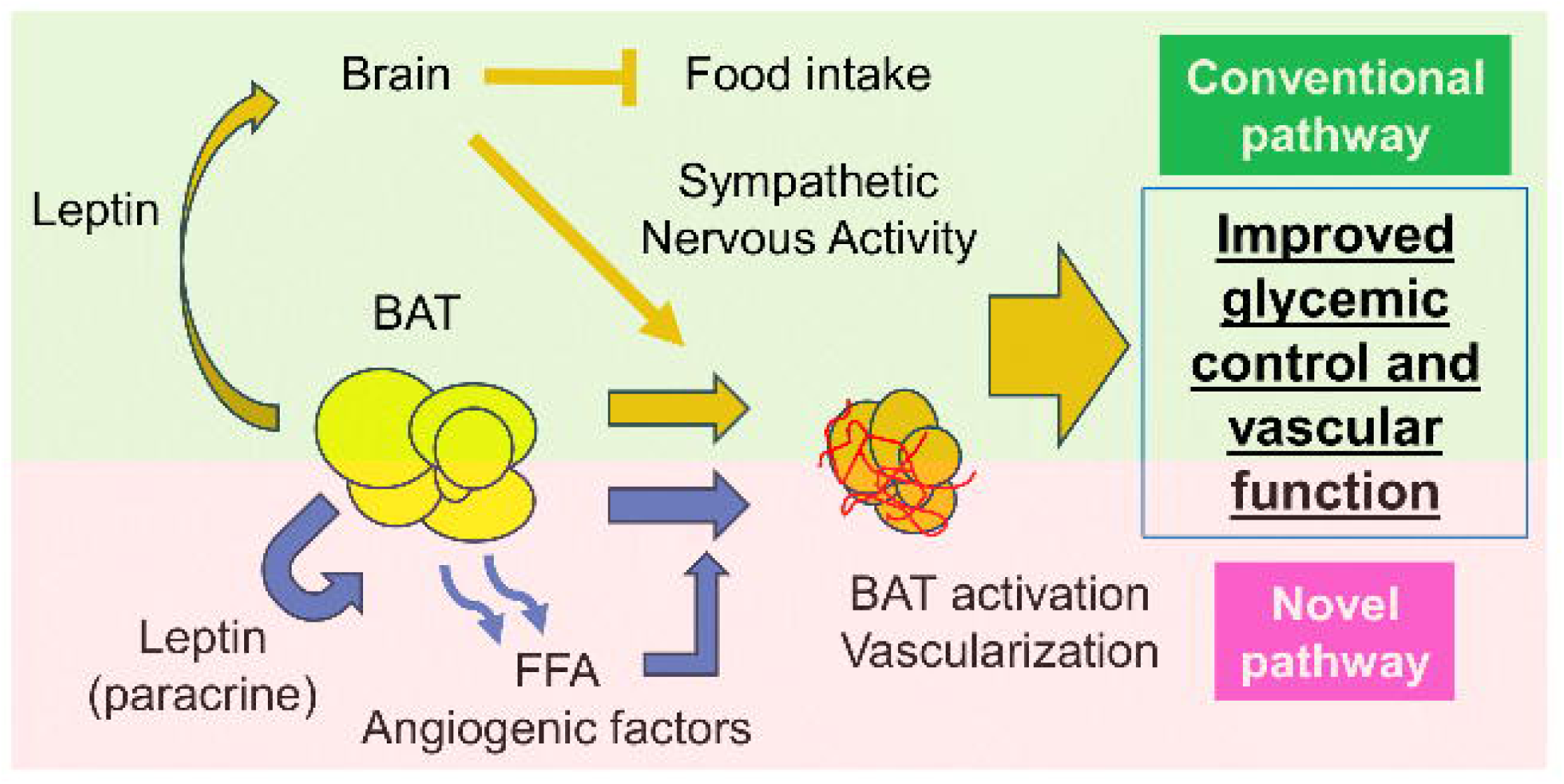

## Introduction

Initially viewed as a satiety hormone (1), the adipokine leptin has rapidly evolved into a prime regulator of energy expenditure and metabolism (2). Interestingly, while leptin receptor (LEPR) is expressed in all metabolic organs, including the adipose tissue (3), leptin control of energy expenditure and metabolic function has primarily been reported to be centrally mediated. Notably, leptin-mediated increases in energy expenditure and lipolysis have been shown to involve central activation of the sympathetic nervous system, which stimulates β-adrenergic receptors on brown (BAT) and white (WAT) adipose tissue to promote the phosphorylation of hormone-sensitive lipase (HSL), the formation of free fatty acids (FFAs), and the activation of uncoupling protein 1 (UCP1) for heat production (4–6). While this mechanism has been reported to be crucial for leptin-mediated increases in energy expenditure and lipolysis, recent *in vitro* evidence, gathered primarily in adipocytes in culture, suggests that leptin could also act directly on adipocytes to stimulate lipolysis (7). Indeed, previous experiments in rat white adipocytes have shown that leptin exposure enhances intracellular triglycerides (TG) hydrolysis and FFA incorporation into TG, ultimately leading to increased net fatty acid efflux (7–10). However, the extent to which the direct effects of leptin on adipocyte metabolism contribute to systemic metabolic regulation and are conserved across all adipose depots, including BAT, in the context of obesity, remains poorly defined. Moreover, it is unclear whether these mechanisms are equally operative in both sexes and whether leptin-induced improvements in adipocyte function confer cardiovascular benefits. To address these questions, we generated and characterized a novel obese mouse model with adipocyte-specific expression of the LEPR, enabling targeted assessment of the metabolic and cardiovascular consequences of adipocyte-restricted leptin signaling in the context of obesity.

## Research Design and Methods

### Models and animals

Male and female obese mice with a selective expression of LEPR in adipocytes were generated by crossing LEPR*^lox^*^TB^ mice (generously provided by Dr. Joel Elmquist at the University of Texas Southwestern Medical Center) with mice expressing Cre recombinase under the control of the adiponectin promoter (APN-Cre mice, Jackson Laboratory, Strain #028020). The LEPR*^lox^*^TB^ model includes a transcriptional blockade cassette inserted after exon 16, disrupting the expression of both the short form (LEPRa), which has limited signaling capacity, and the long signaling-competent form (LEPRb), while leaving the soluble form (LEPRe) intact, as all these isoforms share the same promoter and upstream exons (11). The targeted genetic construct was confirmed by genotyping. Experiments were conducted in 14 to 17-week-old littermates, fed a standard mouse chow (Teklad Global 18% Protein Rodent Diet) and provided water *ad libitum*. LEPR*^lox^*^TB^ x APN-Cre mice were compared to sex and aged-match LEPR*^lox^*^TB^ mice. In addition, wild-type C57BL/6J mice (3–4 months old, both sexes), maintained in our in-house colony, were used for comparison of physiological LEPR expression levels in adipose tissues. All animal investigations were conducted in an American Association for the Accreditation of Laboratory Animal Care-accredited facility with studies approved by the Institutional Animal Care and Use Committee (Protocol #2011-0108). For euthanization procedures, mice were first anesthetized in a closed chamber using 5% isoflurane (1 L/min O_2_) before decapitation. Isolated tissues were snap-frozen in liquid nitrogen before storage at −80°C.

### Indirect calorimetry and body composition

Body composition was determined in a subset of mice using Bruker minispec LF90 TD-Nuclear magnetic resonance (NMR) analyzer. Indirect calorimetry measurements, including oxygen consumption, carbon dioxide respiration, respiratory exchange ratio (RER), heat production, and food and water intake, were measured in plexiglass respiratory chambers using open-circuit Oxymax Comprehensive Lab Animal Monitoring Systems (CLAMS; Columbus Instruments). Non-invasive measurements of the metabolic parameters described earlier were monitored every 5 to 10 minutes for 3 days following a 24-hour acclimation period as previously described (12).

### Infrared Thermography for BAT Activity

Fur over the interscapular region was carefully shaved. Each mouse was then placed in a small cage and allowed to acclimate for 20 minutes. Thermal images were acquired using a FLIR T540 camera, and the interscapular region of interest (ROI) was analyzed using FLIR Tools software (v6.4) to quantify BAT temperature.

### HbA1c, glucose tolerance test and insulin tolerance test

HbA1c was measured at the time of sacrifice in using the A1CNow Self Check system (PTS Diagnostics, Whitestown, IN, USA). Values below the measurable range, shown as <4.0% in the device, were recorded as 3.9%. Intraperitoneal glucose (IPGTTs) and insulin (ITTs) tolerance tests were conducted in conscious fasted mice (16 h fasting for IPGTT; 6 h fasting for ITT). Blood glucose levels was measured at baseline and 15, 30, 60, 90, and 120 minutes after an intraperitoneal bolus injection of glucose (1 g/kg body weight) for IPGTT, and at the same time points plus an additional measurement at 45 minutes after insulin injection (4 U/kg body weight, Humulin R Insulin, 100 units/mL; Eli Lilly, Indianapolis, IN, USA) for ITT.

### Measurements of plasma insulin, leptin, triglycerides and cytokines levels

Plasma leptin (EZML-82K, Millipore Sigma), insulin (EMINS, ThermoFisher Scientific), and triglycerides (TGs, L-Type Triglyceride M Enzyme Color A and B and Multi-Calibrator Lipid, Fujifilm) were measured by ELISA following the manufacturer’s instructions. TGs were also measured in BAT homogenates. Plasma cytokines including TNF-α, IFN-γ, IL-1α, IL-1β, IL-6, IL-10, IL-17A, IL-12p70, GM-CSF, IL-23, IFN-β, MCP-1, and IL-27 were measured using the Mouse Inflammation 13-Plex Panel (FbBA243) as previously described (13).

### Blood pressure measurements

Blood pressure (BP) measurements were obtained using the CODA™ Noninvasive Blood Pressure System (CODA® High Throughput System, KENT Scientific, Torrington, CT, USA) as previously described (13; 14). Mice were trained to the tail-cuff procedure for 3 consecutive days to minimize stress. On the day of measurement, mice were placed in appropriately sized holders on a pre-warmed table in a quiet room (22 ℃), and VPR cuffs were attached to the distal tail. After a 15-min acclimation period, systolic BP was recorded over 25 inflation-deflation cycles per session. Only accepted readings were included in the analysis; sessions with fewer than 10 accepted readings were repeated or omitted. Due to known limitations of tail-cuff measurements for diastolic BP and heart rate (15), only systolic BP values are reported.

### Pulse wave velocity

Pulse wave velocity (PWV) was assessed as an index of aortic stiffness using the INDUS Doppler Flow Velocity System (INDUS Instruments, Webster, TX, USA). Mice were anesthetized and placed supine on a heated platform to maintain body temperature. Doppler probes were positioned over the transverse aortic arch and the abdominal aorta to record flow velocity waveforms at two distinct vascular sites simultaneously. The time delay (Δt) between the foot of the systolic upstroke at the proximal and distal sites was calculated. The distance (Δd) between the probe positions was measured externally using anatomical landmarks. PWV was then determined as the ratio of Δd to Δt (PWV = Δd / Δt). Measurements were synchronized with electrocardiogram signals to ensure precise timing of pulse wave detection.

### Morphological characterization of BAT

Suprascapular BAT was embedded in paraffin and sectioned for hematoxylin and eosin (H&E), DAPI and isolectin B4 (Griffonia Simplicifolia Lectin I, Isolectin B4, 1:100, Biotinylated B-1205-.5, Vector Laboratories) staining and analysis under fluorescence microscopy. For presentation purposes, contrast and brightness of isolectin-stained images were uniformly adjusted using ImageJ. For lipid droplet analysis, 3 to 5 randomly selected fields at 200x magnification were analyzed using ImageJ. Images were converted to grayscale, binarized, and processed with the watershed algorithm to segment individual droplets. Droplets were defined as structures with a black cross-sectional area >20 µm² (diameter >5 µm) and circularity between 0.05 and 1. Non-lipid structures, such as small vessels or artifacts (irregularly shaped rather than oval or circular elements), were manually excluded. For vascular assessment, the area stained by Texas Red was extracted, inverted, binarized, and quantified by ImageJ.

### Vascular Reactivity

Thoracic aortas were dissected surgically from LEPR*^lox^*^TB^ or LEPR*^lox^*^TB^ x APN-Cre mice and cleaned of surrounding fat before being mounted on a wire myograph (DMT, Aarhus, Denmark). Endothelial function was determined according to the protocol previously described (13; 16). Briefly, concentration-response curves to acetylcholine (ACh, A6625 Sigma Aldrich) and sodium nitroprusside (SNP, 567538 Sigma-Aldrich) were conducted in serotonin (5HT, 153-98-0, Cayman Chemical) pre-constricted vessels to assessed endothelium-dependent and independent relaxation. Vascular contractility was also assessed in response to depolarization (Potassium Chloride, KCL, 80mM) and cumulative concentration-response curves to the α_1_-adrenergic receptor agonist phenylephrine (Phe, P6126 Sigma-Aldrich).

### RT-qPCR analysis

A Trizol based method (Qiagen, #15596018, Carlsbad, CA, USA) was employed to extract RNA from abdominal aorta, heart, liver, muscle (iliopsoas), hypothalamus, whole brain without hypothalamus, BAT, subcutaneous adipose tissue (SAT, inguinal), visceral adipose tissue (VAT, perigonadal), and perivascular adipose tissue (PVAT, peri thoracic aorta). Subsequently, complementary DNA library was produced using SuperScript III (ThermoFisher, Waltham, MA). RT-qPCR was performed on the generated cDNA using SYBR™ Select Master Mix (Applied Biosystems™, #4472908, Lithuania) and primers listed in Supplementary Table 1. Ct values were normalized to 18S or GAPDH within the sample (ΔCt), which is subsequently normalized to the control group (ΔΔCt) to generate the relative gene expression (2^-ΔΔCt^). DNA was extracted from BAT using DNeasy Blood & Tissue Kits (Qiagen, #15596018, Carlsbad, CA, USA) and subsequently used to quantify the mitochondrial DNA (mtDNA)-to-nuclear DNA (nuDNA) ratio.

### Western blotting

Mouse tissues were homogenized in RIPA buffer supplemented with protease and phosphatase inhibitors and measured for protein concentration by BCA assay (ThermoFisher), as previously described (17). Tissue homogenates (15 µg) were separated via SDS-PAGE and transferred to Immobilin-P poly(vinylidene fluoride) membranes. Immunoblots were probed with antibodies for CD31 (Abcam, ab124432, 1:500), UCP1 (Abcam, ab23841, 1:1000), Anti-β-Actin-Peroxidase antibody (Sigma-Aldrich, A3854, 1:1000), HSL (Cell Signaling, #4107, 1:1000), phospho-HSL (Ser563, Cell Signaling, #4139, 1:1000), ATGL (30A4 ,Cell Signaling, #2439, 1:1000), phospho-ATGL (S406, Abcam, ab317610, 1:1000), TOM20 (Cell Signaling, #42406, 1:1000), LEPR (Abcam, ab104403, 1:1000), vinculin (Cell Signaling, #13901, 1:1000), AKT (Cell Signaling, #9272, 1:1000), and pAKT (Thr308, Cell Signaling, #9275, 1:1000) used with 1% bovine serum albumin (BSA, Millipore Sigma) and TRIS-buffered saline (TBS, 10x) pH 7.4 with MilliQ and 0.05% of tween-20 (Fisher BioReagents) for 1 hour at room temperature. Mouse (goat anti-mouse IgG-HRP, Bio-Rad #1706516, 1:1000) and rabbit (goat anti-rabbit IgG-HRP, Bio-Rad #1705046, 1:5000) secondary antibodies diluted by 1% BSA TBS-T were employed. Membranes were revealed using the ChemiDoc Imaging System (Bio-Rad) Version 3.0.1.14.

### Statistics

All data are presented as mean ± SEM. Whenever applicable, un-paired two-sample t-test, one-way analysis of variance (ANOVA) followed by Dunnett post hoc test, or two-way ANOVA followed by Sidak’s post-hoc test were used to analyze the data (GraphPad Prism 10; GraphPad Software Inc., La Jolla, CA). P-values or multiple-comparison adjusted p-values < 0.05 were considered significant.

### Data and Resource Availability

Supporting data are available from the corresponding authors upon reasonable request.

## Results

### Selective restoration of adipocyte LEPR improves BAT structure and function in male mice only

To confirm selective adipocyte LEPR restoration in LEPR*^lox^*^TB^ x APN-Cre, LEPR transcript levels were quantified by qRT-PCR in the heart, liver, muscle, hypothalamus, brain without hypothalamus, BAT, SAT, VAT, and PVAT. As shown in Supplementary Fig. 1A and 2A, male and female LEPR*^lox^*^TB^ x APN-Cre mice expressed the long, LEPRb, and short form, LEPRa, of the leptin receptor in all fat depots but not in other organs, confirming the selectivity of our mouse model. Re-expression of the LEPR in LEPR*^lox^*^TB^ x APN-Cre mice had only minor effects on the circulating form, LEPRe, (Supplementary Fig. 1C and 2C), whose expression was mostly unaffected, with the exception of the male SAT that exhibited a significant increase, and the female liver, muscle, and BAT that showed significant reductions. Consistent with the transcript data, BAT LEPRb protein expression was significantly increased in both sexes (Supplementary Fig. 3A). Interestingly, male LEPR*^lox^*^TB^ x APN-Cre mice exhibit a 4-fold and 62-fold increase in BAT LEPRb and LEPRa expression, respectively, compared to female counterparts. Other adipose tissues did not exhibit sex differences in either LEPRb or LEPRa expression (Supplementary Fig. 3B). Analysis of inter-adipose tissue differences in both males and females revealed that although LEPRb expression was restored to a similar extent in BAT, VAT and PVAT, only SAT exhibited a higher restoration in expression in comparison to BAT in both sexes. However, LEPRa expression did not differ between adipose depots in both males and females (Supplementary Fig. 3C). To assess the level of LEPR re-expression, obese LEPR*^lox^*^TB^ x APN-Cre mice were compared to lean wild-type mice. Male and female LEPR*^lox^*^TB^ x APN-Cre mice exhibit approximatively 14-fold and 6-fold higher BAT LEPRb, respectively, compared to lean wild-type C57Bl/6 mice but similar BAT LEPRa levels (Supplementary Fig. 3D). Lastly, lean male and female wild-type C57Bl/6 mice present with no differences in BAT LEPRb and LEPRa levels (Supplementary Fig. 3D).

In male mice, selective restoration of adipocyte LEPR led to a significant increase in BAT weight with no alteration in other fat pad weights, total body weight, or total fat volume (Fig. 1A and B). In females, restoration of adipose tissue LEPR selectively reduced SAT and increased the percentage of fluid volume (Fig. 1A and B). These minor changes in body composition were observed without changes in circulating leptin levels (Fig. 1C), food intake, fluid consumption, and activity, in either male or female mice (Supplementary Fig. 4A and B). BAT is a central regulator of energy expenditure whose weight correlates positively with its activity (18). Therefore, we employed indirect calorimetry to assess the effects of adipocyte LEPR re-expression on energy expenditure. Selective expression of LEPR in adipose tissue increased oxygen consumption (VO_2_), carbon dioxide production (VCO_2_), and heat production, in male mice only, reflecting an increase in energy expenditure (Fig. 1D). This increase in heat production was mainly localized in BAT, as demonstrated by infrared thermography (Fig. 1E). However, LEPR re-expression did not alter RER.

**Figure 1.**
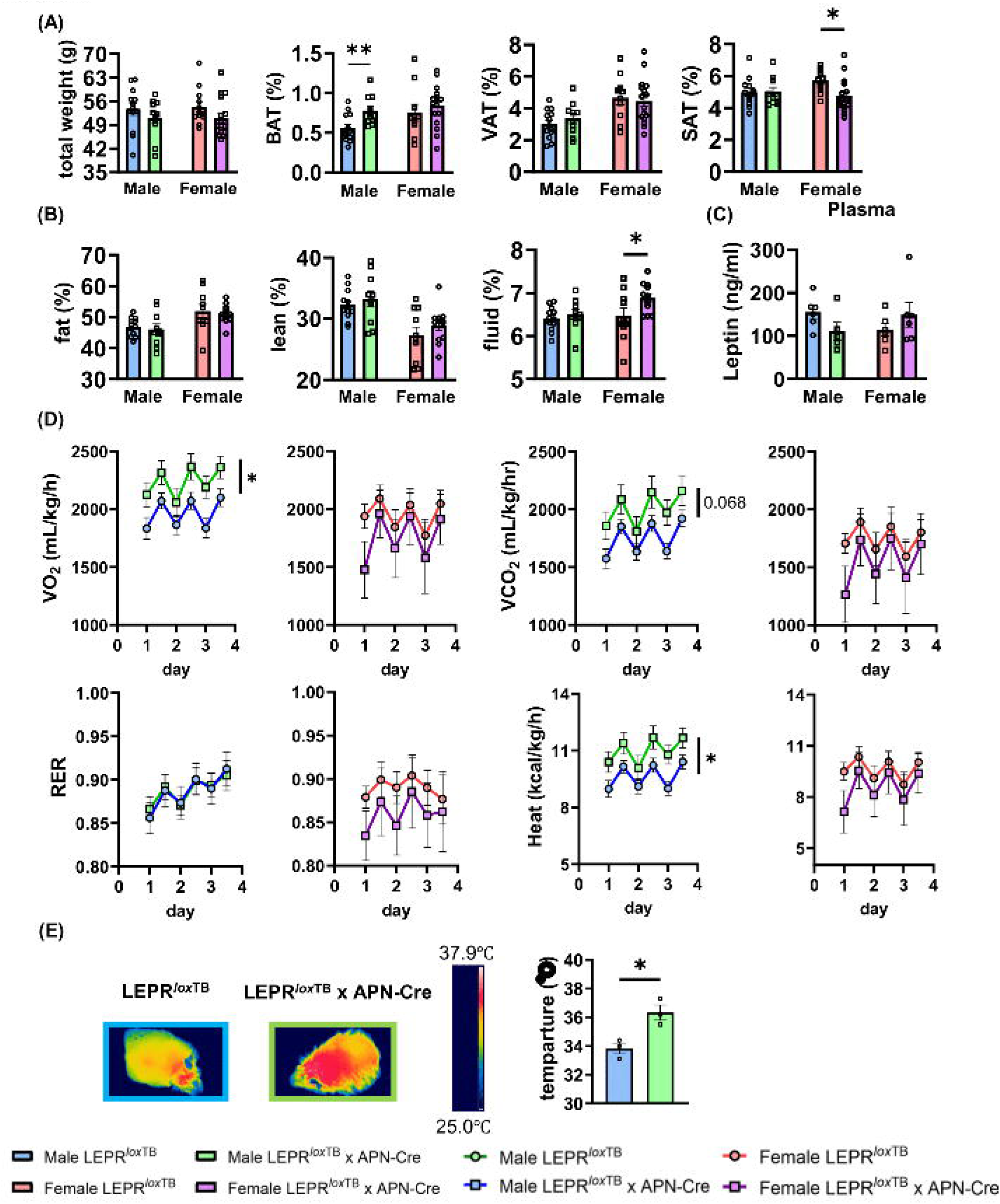
Restoration of adipocyte LEPR enhanced BAT weight and thermogenic activity in male mice only. (A, B) Body and tissue weights, expressed as percentage of total body weight, measured by NMR at sacrifice. (C) Plasma leptin levels. (D) Measurements of VO₂, VCO₂, and heat production in LEPR*^lox^*^TB^ and LEPR*^lox^*^TB^ x APN-Cre male and female mice, conducted via indirect calorimetry. (E) The temperature of the interscapular BAT region measured by infrared thermography in male LEPR*^lox^*^TB^ and LEPR*^lox^*^TB^ x APN-Cre mice. Blue, green, red, and purple bars or lines represent male LEPR*^lox^*^TB^, male LEPR*^lox^*^TB^ x APN-Cre, female LEPR*^lox^*^TB^, and female LEPR*^lox^*^TB^ x APN-Cre, respectively. Data are presented as mean±SEM, n=3-16. *P < 0.05, **P < 0.01.

Based on the selective effects of LEPR restoration on BAT size and function, we investigated BAT morphology. Adipose tissue LEPR restoration decreased BAT adipocyte size and increased adipocyte density in males (Fig. 2A) but not in females (Supplementary Fig. 5A). We also employed qRT-PCR to quantify genes related to browning, including UCP1, PRDM16, and NFIA (19). Adipocyte LEPR restoration increased UCP1, PRDM16, NFIA, and PPARGC1A expression in male (Fig. 2B) but not in female mice (Supplementary Fig. 5B), supporting an increase in browning in males only. DIO2, TH, and ADRB3 expression remained unchanged, indicating that sympathetic activation or DIO2-mediated mechanisms are unlikely to account for the increase in UCP1. (Fig. 2B). Consistent with the transcript data, UCP1 expression was also elevated at the protein level in male LEPR*^lox^*^TB^ x APN-Cre (Fig. 2C). TOM20, a protein marker of mitochondrial mass, was significantly increased in LEPR re-expressing male BAT. However, mtDNA/nuDNA was not significantly altered (Supplementary Fig. 6A), suggesting an increase in mitochondrial function rather than an increase in mitochondrial number in male LEPR*^lox^*^TB^ x APN-Cre mice. Consistent with these observations, pSTAT3 activation was significantly enhanced in male BAT by the restoration of adipocyte LEPR but remained unchanged in female BAT (Fig. 2D, Supplementary Fig. 6B). LEPR restoration led to no increase in browning markers in SAT, VAT, and PVAT (Supplementary Fig. 6C), suggesting primarily BAT-specific effects. However, adipocyte LEPR restoration induced a marked decrease in browning markers (UCP1, NFIA) in female PVAT. As lipolysis increases UCP1 activity and browning (20), we measured circulating and BAT TG levels. We also assessed the degree of activation of adipose triglycerides lipase (ATGL) and HSL activity by evaluating their level of phosphorylation. As reported in Fig. 2E and supplementary Fig. 6D, re-expression of adipocyte LEPR reduced BAT TG levels without altering plasma TG in male mice only. Furthermore, re-expression of adipocyte LEPR significantly increased HSL activity in males (Fig. 2F), which supports an increase in lipolysis in male BAT.

**Figure 2.**
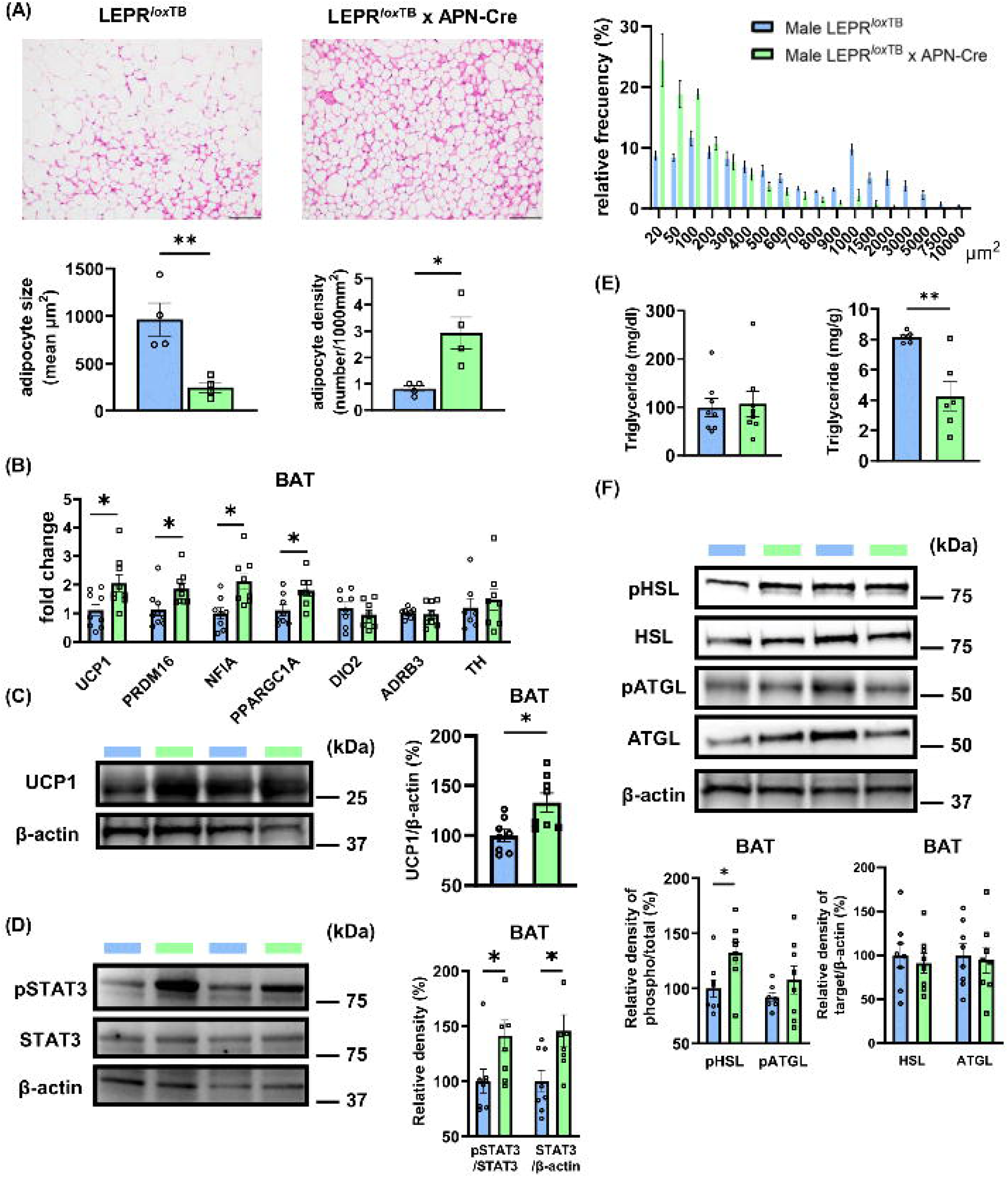
Restoration of adipocyte LEPR induces browning in male BAT. (A) Representative section of BAT from male LEPR*^lox^*^TB^ and LEPR*^lox^*^TB^ x APN-Cre mice stained with H&E staining from which adipocyte size, density and frequency of distribution were measured. Scale bars indicate 50 μ m (B) qPCR quantification of the browning markers, UCP1, PRDM16, and NFIA in male BAT. (C) Representative western blot and quantification of UCP1 in male BAT. (D) Representative western blots and quantification of total and phosphorylated STAT3. (E) Plasma and BAT triglyceride levels in males. (F) Representative western blots and quantification of total and phosphorylated HSL and ATGL. Blue and green bars represent male LEPR*^lox^*^TB^ and male LEPR*^lox^*^TB^ x APN-Cre, respectively. Data are presented as mean±SEM. n=4-9. *P < 0.05, **P < 0.01.

### Selective restoration of adipocyte LEPR increases BAT vascularization

Evidence indicates that BAT perfusion and vascular density contribute to the regulation of its activity (21). Therefore, we measured indices of vascular density in LEPR*^lox^*^TB^ x APN-Cre mice. We quantified markers of endothelial cell numbers, including isolectin B4, CD31, and CDH5, and markers of angiogenesis: VEGFA, HIF-1, HIF-2, ANGPT1, ANGPT2, and TGFβ. All these markers were increased in male LEPR*^lox^*^TB^ x APN-Cre mice compared to LEPR*^lox^*^TB^, supporting higher vascular density and increased angiogenesis in the BAT of male mice (Fig. 3A-C). LEPR re-expression did not lead to an increase in markers of angiogenesis in female LEPR*^lox^*^TB^ x APN-Cre mice (Supplementary Fig. 7).

**Figure 3.**
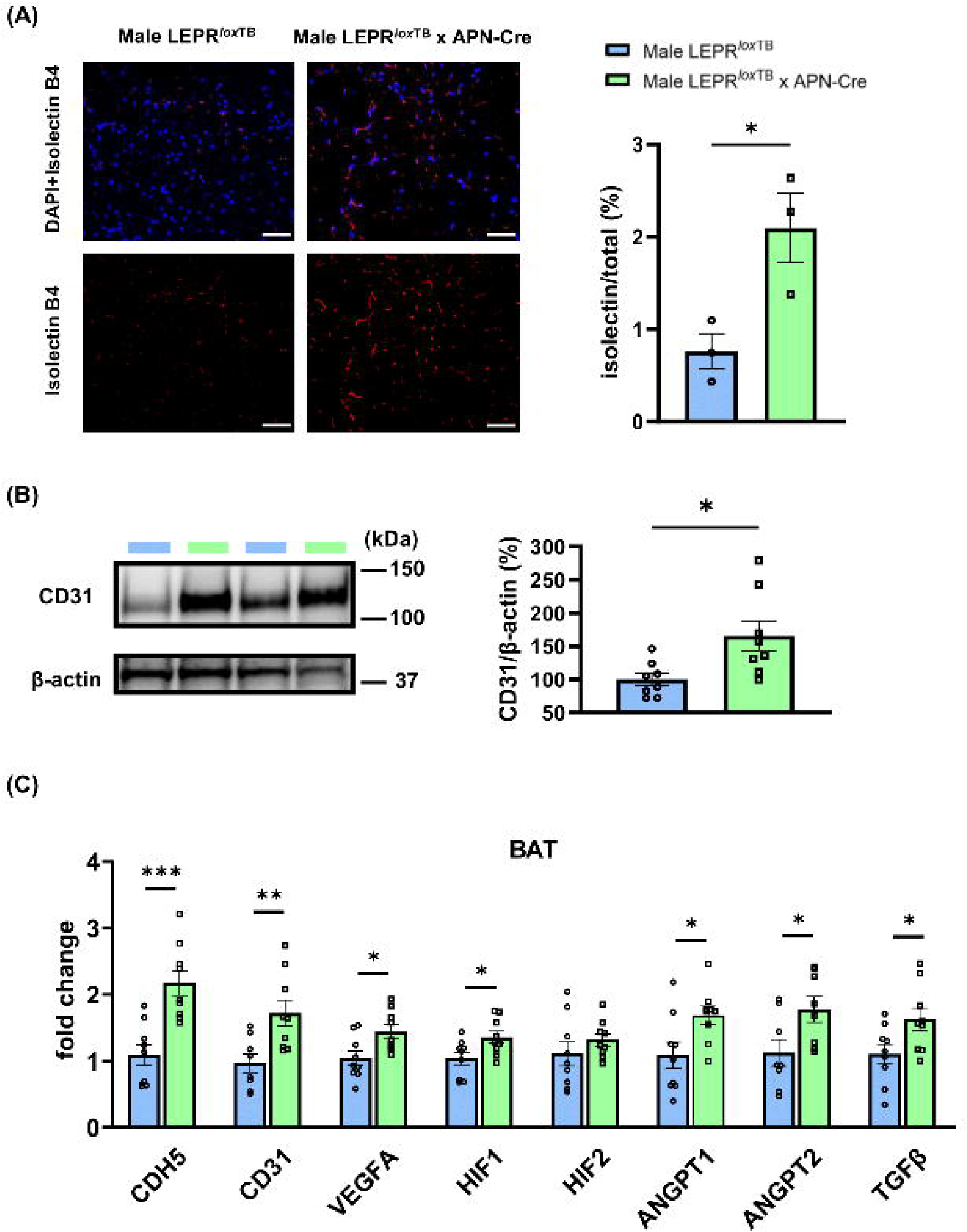
Restoration of adipocyte LEPR enhanced angiogenesis and vascular stabilization. (A) Representative sections of BAT from LEPR*^lox^*^TB^ and LEPR*^lox^*^TB^ x APN-Cre mice stained for isolectin B4 and quantified. Data are presented as the ratio of isolectin B4/total area. Scale bars indicate 60 μm. (B) Representative western blots and quantification of BAT CD31 protein levels. (C) qPCR quantification of endothelial and angiogenesis markers. Blue and green bars represent male LEPR*^lox^*^TB^ and male LEPR*^lox^*^TB^ x APN-Cre, respectively. Data are presented as mean±SEM. n=3-9. *P < 0.05, **P < 0.01, ***P < 0.001.

### LEPR restoration in adipocytes improved glycemic control

BAT plays a key role in systemic glycemic control. Therefore, we assessed glucose and insulin sensitivity in LEPR*^lox^*^TB^ and LEPR*^lox^*^TB^ x APN-Cre. Restoration of adipocyte LEPR significantly lowered HbA1c and plasma insulin levels in male (Fig. 4A and B) but not female mice (Supplementary Fig. 8A and B). Glucose clearance during IPGTT was not markedly improved in either sex (Fig. 4C, Supplementary Fig. 8C). However, a significantly improved insulin response was observed in male LEPR*^lox^*^TB^ x APN-Cre mice only (Fig. 4D, Supplementary Fig. 8D). Enhanced glycemic control in male LEPR-restored mice was supported by increased pAKT/AKT in BAT. In contrast, no changes were observed in muscle pAKT/AKT or in GLUT4 expression in adipose tissue and muscle (Fig. 4E and F).

**Figure 4.**
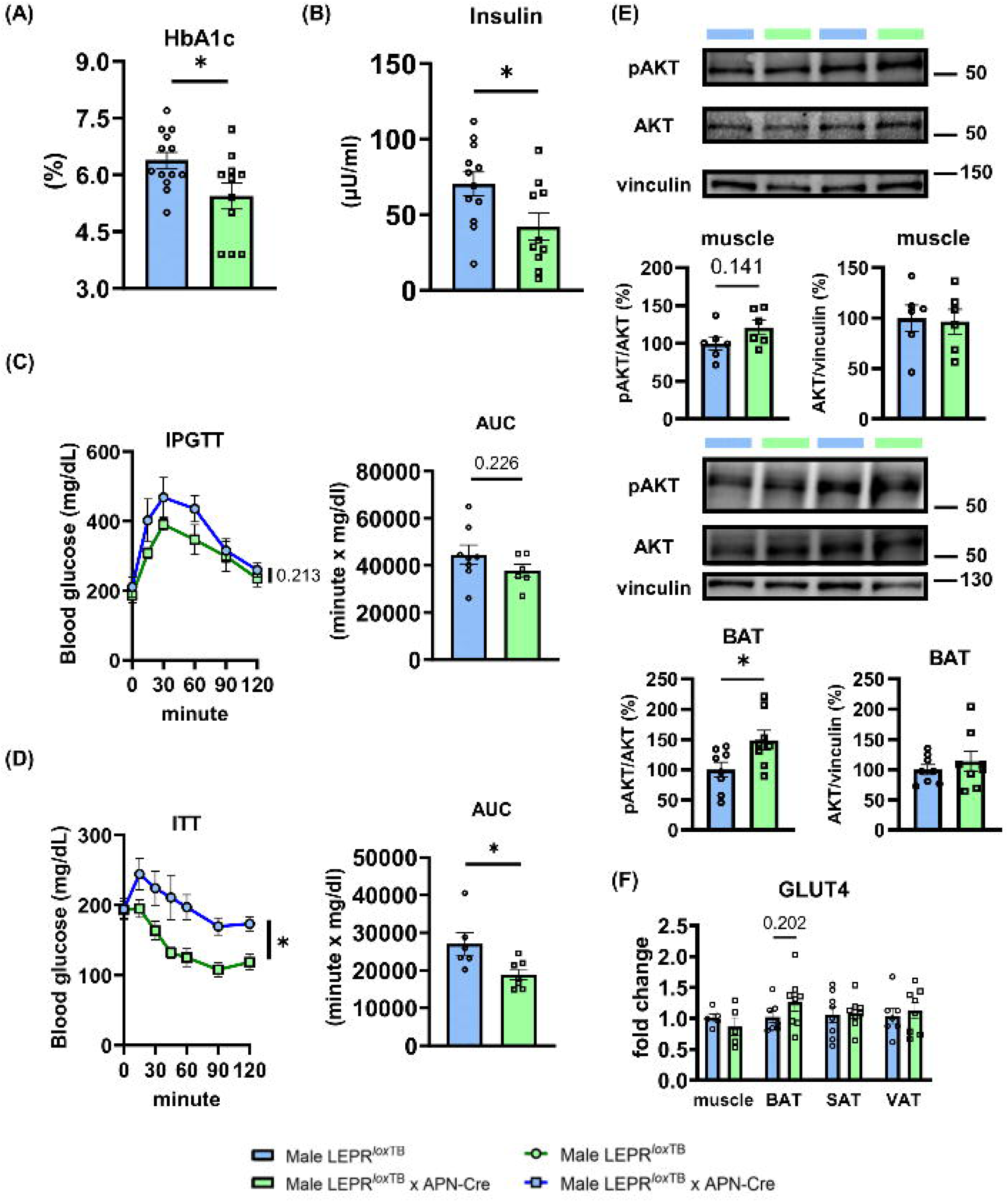
Restoration of adipocyte LEPR improved insulin sensitivity and glucose metabolism in male mice. (A) HbA1c and, (B) Plasma Insulin levels, (C) Blood glucose response to a glucose tolerance test (IPGTT) and area under the curve quantification, (D) Blood glucose response to an insulin tolerance test (ITT) and area under the curve quantification. (E) Representative western blots and quantification of total and phosphorylated AKT. (F) qPCR quantification of GLUT4 in insulin-sensitive tissues such as muscle, BAT, SAT, and VAT. Blue and green bars represent male LEPR*^lox^*^TB^ and male LEPR*^lox^*^TB^ x APN-Cre, respectively. Data are presented as mean±SEM, n=5-14. *P < 0.05.

### LEPR restoration in adipocytes improves cardiovascular function

Consistent with the improved glycemic control, systolic BP (SBP) and arterial stiffness (PWV) were significantly reduced in LEPR*^lox^*^TB^ x APN-Cre males (Fig. 5A and B). Furthermore, restoration of adipocyte LEPR significantly improved endothelium-dependent relaxation assessed by acetylcholine-mediated dilatation without altering endothelium-independent relaxation measured in response to sodium nitroprusside in male mice (Fig. 5C and D). However, in female mice, adipocyte LEPR re-expression did not improve SBP, PWV, or endothelial function (Supplementary Fig. 9A-D). Vascular contractility to depolarization (KCl) and α_1_-adrenergic receptor activation (phenylephrine), remained intact in both male and female LEPR*^lox^*^TB^ x APN-Cre mice (Fig. 5E and F and Supplementary Fig. 9E and F).

**Figure 5.**
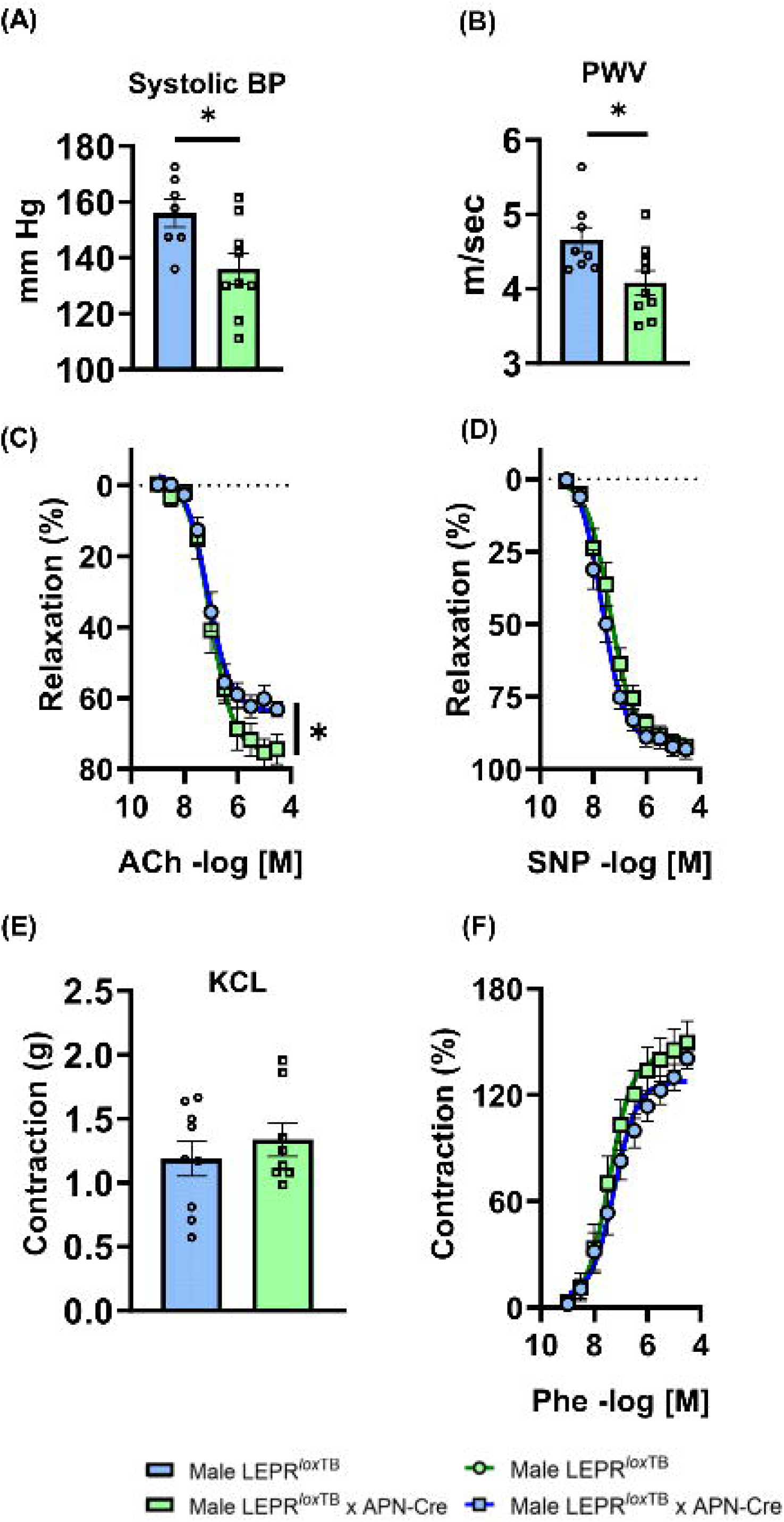
Restoration of adipocyte LEPR improved cardiovascular function including blood pressure, arterial stiffness and endothelial function in male mice. (A) Systolic BP measured by tail-cuff method, (B) Pulse wave velocity (PWV), (C to F) Wire myograph analysis of aorta (C) endothelium-dependent, (D) endothelium independent relaxation, (E, F) constriction to potassium chloride (80mM) and phenylephrine. Blue and green bars represent male LEPR*^lox^*^TB^ and male LEPR*^lox^*^TB^ x APN-Cre, respectively. Data are presented as mean±SEM, n=5-11. *P < 0.05.

### LEPR restoration in adipocytes suppresses cardiovascular inflammation

To assess the extent of the beneficial effects of the improved metabolic function in LEPR*^lox^*^TB^ x APN-Cre, we quantified circulating and tissue markers of inflammation. Restoration of adipocyte LEPR significantly reduced plasma TNF-α, IL-1β, and IFN-γ and led to a trend towards a reduction in IL-17A in male mice (Fig. 6A). This reduction in circulating pro-inflammatory cytokines in male LEPR*^lox^*^TB^ x APN-Cre was associated with a decrease in vascular (aorta) and cardiac markers of oxidative stress and inflammation (NOX2, NOX4, TNF-α), immune cell adhesion (VCAM1, ICAM1), macrophage infiltration (F4/80) and of marker of M1 macrophage activation (iNOS/CD206) (Fig. 6B and C). Restoration of adipocyte LEPR in female mice did not improve plasma or tissue markers of inflammation (Supplementary Fig. 10A, B, and C). The ameliorated systemic and local inflammation observed exclusively in male LEPR*^lox^*^TB^ x APN-Cre mice was associated with increases in the expression of BAT-derived cardioprotective batokines (22; 23) including fibroblast growth factor 21 (FGF21) and neuregulin 4 (NRG4, Fig. 6D). No alterations in BAT FGF21 and NRG4 were reported in female LEPR*^lox^*^TB^ x APN-Cre mice (Supplementary Fig. 10D).

**Figure 6.**
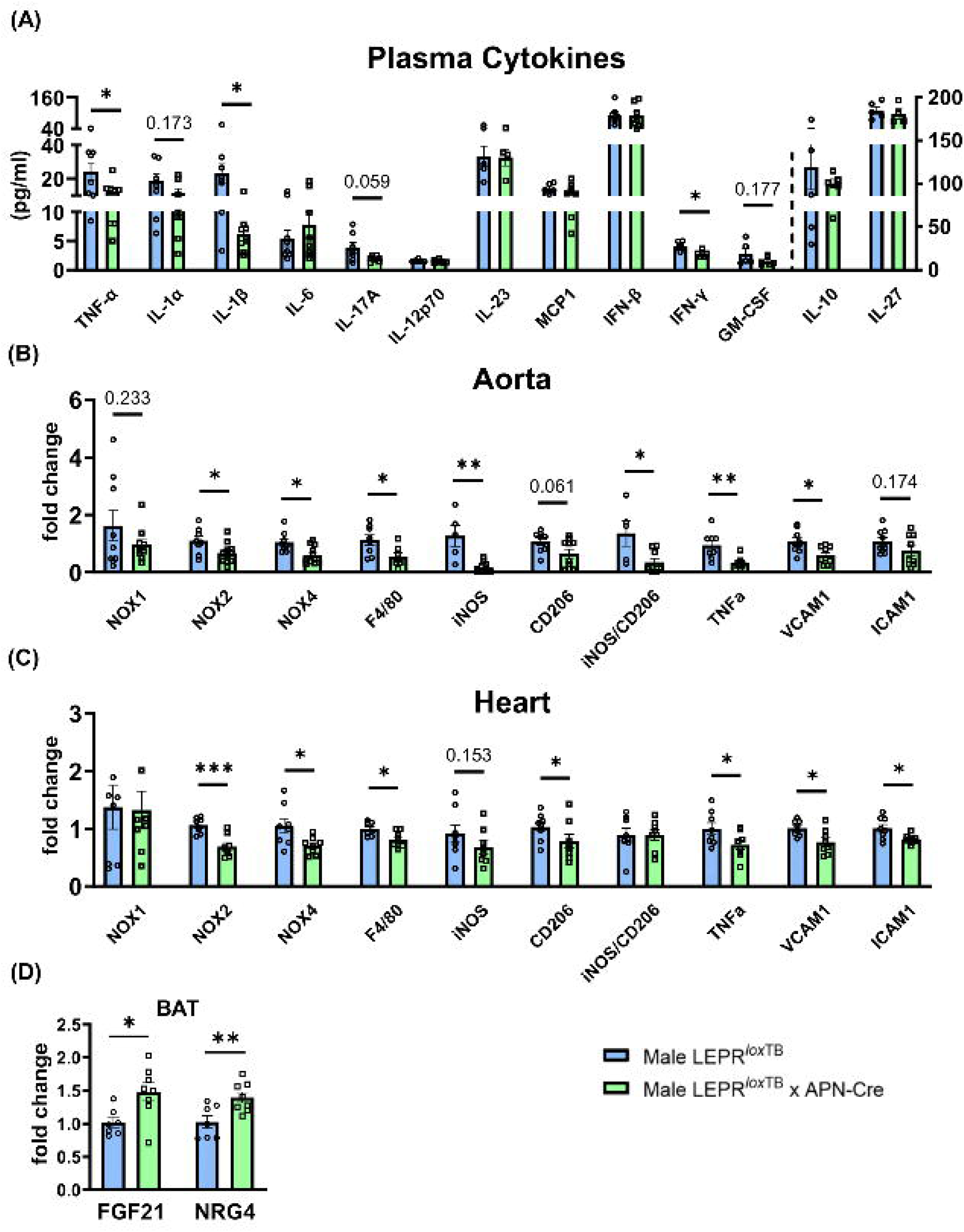
Restoration of adipocyte LEPR reduced systemic and tissue inflammation in male mice. (A) Plasma cytokine, (B) Aorta and (C) Heart quantification of local inflammation via qPCR analysis. (D) qPCR quantification of FGF21 and NRG4 in BAT. Blue and green bars represent male LEPR*^lox^*^TB^ and male LEPR*^lox^*^TB^ x APN-Cre, respectively. Data are presented as mean±SEM, n=4-12. *P < 0.05, **P < 0.01, ***P < 0.001.

## Discussion

To test the potential direct role of adipose leptin signaling in controlling energy expenditure and metabolism, we generated an obese mouse with a selective expression of LEPR in adipocytes. While characterizing the phenotype of these animals, we reported no significant alterations in body composition but a significant increase in male BAT mass accompanied by an increase in energy expenditure, a reduction in BAT adipocyte size, a higher BAT adipocyte density, an enhanced lipolysis and an elevated BAT vascular density in male mice only. Remarkably, the male specific improved BAT structure and function were associated with ameliorated insulin sensitivity and HbA1c, which likely contributed to an ameliorated cardiovascular function reflected by a decreased blood pressure, a reduced arterial stiffness, and an enhanced endothelium-dependent relaxation in males. Relevant to these findings are the mouse model of selective adipose leptin signaling, the selective effects of leptin on BAT, the sex specificity of the mechanism, and the cardiovascular benefits of restoring adipose leptin signaling.

To evaluate the specific effect of adipocyte leptin signaling on metabolism, we took advantage of the conditional LEPR knockout mice in which LEPR can be restored in tissues of interest. Using this model and an adipocyte Cre line, we restored LEPRb and LEPRa expression exclusively in adipose depots of both male and female mice. Adipose LEPRa levels have been restored to levels matching those of lean sex-matched wild-type control mice, while the only known functional and signaling receptor, LEPRb, was re-expressed to levels 6-fold and 14-fold higher, respectively, in female and male mice, than in the BAT of lean wild-type animals. These levels may appear supraphysiological. However, as the effects of obesity on BAT LEPRb levels are unknown, a comparison with an alternative model of obesity would provide a clearer assessment of their pathophysiological relevance. Nevertheless, this potential limitation does not diminish the significance of the model, as LEPR*^lox^*^TB^ x APN-Cre mice offer a unique opportunity to delineate the specific contribution of adipocyte leptin signaling in the setting of severe obesity and diabetes, and importantly, in the absence of central leptin action. Given the limited knowledge of the function of LEPRa, which lacks intracellular signaling (11), all the effects reported in the present study have been attributed to LEPRb.

Remarkably, while LEPR*^lox^*^TB^ x APN-Cre mice exhibit a restoration of LEPR in all adipose depots, enhanced thermogenic activity was reported exclusively in the BAT, and more specifically in male BAT. These findings are consistent with Pereira’s observations, showing, in the exact same mouse model, that adipocyte LEPR restoration did not alter white adipose tissue (WAT) function, but in contradiction with their findings showing no increase in male BAT UCP1 levels (24). The older age of their mice and the relatively superficial analysis of the BAT function could potentially be the cause of the discrepancy between the 2 studies. Nevertheless, these 2 studies reporting no effects of LEPR restoration on WAT beiging and lipolysis tend to support a limited physiological role for the leptin-mediated lipolysis reported in rodent visceral and subcutaneous adipocytes in culture (7–10). *In vitro* experiments in children and adult human adipocytes in culture further minimize the role of leptin in WAT lipolysis (25). Intriguingly, these findings raise the question of the origin of the selective effects of leptin on BAT. No differences in LEPR expression were observed between BAT, VAT, and PVAT in male and female LEPR*^lox^*^TB^ x APN-Cre mice, only SAT exhibited a marked increase in LEPR expression, in comparison to BAT in both sexes. Therefore, these data rules out LEPR expression level as an explanation for the tissue specificity. The unique nature of BAT cells, which originate from myogenic factor 5 (Myf5) positive progenitor cells, whereas WAT adipocytes derive from Myf5-negative progenitor cells (26; 27), may contribute to the distinct response to leptin. Similarly, activation of different signaling pathways within white and brown adipocytes, as well as the potential development of a tissue-specific resistance to leptin, may explain the differences. Based on the very high circulating leptin levels in LEPR*^lox^*^TB^ and LEPR*^lox^*^TB^ x APN-Cre mice, one can reasonably speculate that the BAT remains leptin sensitive while WAT may have developed resistance to leptin. However, addressing these questions is out of the scope of the present study.

Our study revealed for the first time, *in vivo* and in pathophysiological conditions, a direct role for leptin in the control of BAT function and shows that BAT leptin signaling increases lipolysis. BAT function is critically reliant on UCP1 expression, which was upregulated following restoration of LEPR expression. While the precise mechanisms by which adipocyte-specific leptin signaling regulates UCP1 levels remain to be elucidated, our findings suggest several potential pathways that may underlie this regulation. Leptin classically activates STAT3 signaling (11). However, although we report increased BAT STAT3 activation, recent evidence suggests that STAT3 inhibition can paradoxically enhance UCP1 expression in brown adipocytes (28). This likely indicates that other leptin-activated pathways such as AKT, AMPK, or MAPK may contribute to the induction of UCP1 and thermogenic genes (29–33). Interestingly, BAT LEPR restoration increased pAKT/AKT suggesting that this pathway could be involved. Other mediators known to enhance UCP1, such as cAMP and PPARγ, may also contribute to leptin-mediated UCP1 expression, as leptin has been shown to increase PPARγ in endothelial cells and activate cAMP signaling in adipose tissue macrophages (34–36). As UCP1 exerts its thermogenic function within the inner mitochondrial membrane (37), we also quantified markers of mitochondria function and abondance. BAT LEPR restoration led to increases in BAT PGC1α, a master regulator of mitochondrial number, function, and quality (38), as well as TOM20, a marker of mitochondrial content, but without altering the mtDNA/nuDNA ratio. These finding suggests that the enhanced thermogenic phenotype is likely driven primarily by improved mitochondrial function rather than by a substantial increase in mitochondrial abondance. Given the significant induction of UCP1 and PGC1α, it is plausible that leptin signaling promotes mitochondrial activity or remodeling rather than large-scale biogenesis. However, further studies are required to identify the exact molecular mechanisms whereby leptin controls UCP1 levels and BAT activity.

Our results also imply increased free fatty acid production through increases in HSL activity and increase in BAT vascularization. Indeed, while BAT activity has been positively related to BAT vascular density (21; 39), we showed that LEPR restoration significantly increased BAT capillary density, as shown by increases in isolectin B4 staining, and endothelial markers including CD31 and CDH5. Moreover, consistent with the central role of VEGFA in the control of BAT vascularization (21; 40-42), we reported increased levels of VEGFA in LEPR*^lox^*^TB^ x APN-Cre mice BAT. VEGFA is produced by adipocytes in adipose tissue (43). Consistent with the literature (44), we showed that restoration of leptin signaling activates the STAT3 pathway in adipocytes, which is known to upregulate VEGFA in retinal endothelial cells (45). Given that adipocytes can produce VEGFA, it is plausible that leptin signaling directly enhances VEGFA expression in BAT adipocytes as well, promoting angiogenesis. In addition, our study showed that other angiogenic factors, including ANGPT1, ANGPT2, TGF-β, and HIF1, were also upregulated following adipocyte LEPR restoration. ANGPT2 has been reported to be directly enhanced by leptin in adipocytes (46), and could act synergistically with VEGFA to promote endothelial cell proliferation and vessel maturation (47). TGF-β and HIF1 further enhance the angiogenic response by stabilizing new capillary networks and adapting BAT to increased metabolic demands, the former of which is related to brown and beige adipocyte differentiation (48–50). These findings suggest that leptin signaling in adipocytes orchestrates a complex angiogenic program beyond just VEGFA activation.

Another major finding of our study is the observation that LEPR-mediated BAT remodeling and activation was exclusively observed in males. The higher restoration of LEPR in males’ BAT compared to females would be the easiest explanation. However, the obesity consecutive to LEPR deficiency did not alter BAT morphology and function in females, and LEPR restoration in male BAT did in fact abolish the sex difference. This support sex differences in the control of BAT tissue morphology and function and suggest leptin-independent mechanisms in females. The preserved BAT function and morphology in obese females is likely thanks to estrogen which drives BAT remodeling and increases energy expenditure through peripheral estrogen receptor α signaling (51; 52). Thus, it is likely that BAT morphology in females had already been optimized by estrogen, preventing further structural adaptation upon adipocyte-specific LEPR restoration.

Added to the enhanced BAT function, adipocyte LEPR restoration markedly improved cardiovascular function in male mice as reflected by a reduction in blood pressure and arterial stiffness and an improved endothelial function. Given the inherent limitations of the tail-cuff technique, including its susceptibility to stress-induced variability and substantial operator dependence, the absolute blood pressure values obtained should be interpreted with caution. However, the reduced blood pressure reported is corroborated by consistent changes across multiple independent cardiovascular parameters, collectively supporting a genuine decrease in blood pressure. This improved cardiovascular function was accompanied by lower systemic and tissue inflammation. Sustained hyperglycemia and insulin resistance, as seen in diabetes, are a major cause of cardiovascular complications including hypertension, atherosclerosis, inflammation and endothelial dysfunction (53–57). Furthermore, enhanced energy expenditure reduces the risk for cardiovascular diseases via improved insulin sensitivity (58). Therefore, one can reasonably speculate that LEPR-mediated BAT activation alleviates the deleterious effects of hyperglycemia on cardiovascular function by the improvement of insulin sensitivity. In addition to improving glycemia and reducing vascular inflammation, BAT can influence cardiovascular function via the secretion of endocrine factors such as FGF21 and NRG4, known as cardioprotective batokines (22; 23). We observed that adipocyte LEPR restoration increased BAT FGF21 and NRG4 transcript levels exclusively in males. Remarkably, females that did not demonstrate improved cardiovascular function with adipocyte LEPR re-expression, exhibited no alterations in batokine transcript levels. While these data are only correlative, they supports a potential contribution of batokines to the observed improvements in blood pressure and vascular function in males. Furthermore, these latter data substantiate the concept that BAT leptin signaling controls glycemia, and cardiovascular function in males only. Indeed, we reported that irrespective of the presence of adipocyte LEPR, obese female mice exhibited no alteration in BAT function and glycemia but surprisingly developed cardiovascular alterations arguing for a dissociation between adipocyte leptin signaling, BAT thermogenic capacity and cardiovascular health in females.

In conclusion, we report for the first time a male-specific role of adipocyte leptin signaling in the regulation of brown adipose tissue function, particularly in the modulation of BAT lipolysis and vascularization. These processes, in turn, contribute to the regulation of systemic glycolysis and cardiovascular function. Our findings highlight adipocyte leptin signaling as a potential therapeutic target for the treatment of metabolic and cardiovascular disorders in males.

## Acknowledgements

We thank the Augusta University Medical College of Georgia Electron Microscopy & Histology Core Facility (RRID:SCR_026810) for assistance with histological analyses. We appreciate the cooperation of the following individuals for their technical support: Jing Zhao (Electron Microscopy & Histology Core, Medical College of Georgia, Augusta University) for H&E staining and immunostaining; James Mints (Vascular Biology Center, Medical College of Georgia, Augusta University) for his help with NMR and CLAMS measurements. We also thank FLIR Systems, Inc. for providing the FLIR T540 thermal imaging system used in this study. Dr. Eric J. Belin de Chantemèle is the guarantor of this work and, as such, had full access to all the data in the study and takes responsibility for the integrity of the data and the accuracy of the data analysis

## Author contributions

The authors’ contribution for this paper is as follows; Yoichi Ono: Conceptualization, Methodology, Formal analysis, Data Curation, Writing - Original Draft, Figures preparation, Funding acquisition. Simone Kennard, Ben Wall, Jing Ma: Mouse Management, Material Procurement. Eric J Belin de Chantemèle: Conceptualization, Data Curation, Supervision, Project administration, Writing - Original Draft, Funding acquisition.

## Data and resource availability statements

The datasets and resources generated and analyzed in the current study are available from the corresponding author upon reasonable request.

