## Supplemental Material for "Adipocyte Leptin Signaling Regulates Glycemia and Cardiovascular Function via Enhancing Brown Adipose Tissue Thermogenesis in Obese Male Mice"

**Running head**: Direct leptin control of brown adipose tissue

*Corresponding Authors: Eric J. Belin de Chantemèle, DSc. FAHA
Vascular Biology Center
Department of Medicine (Cardiology),
Medical College of Georgia at Augusta University
1460 Laney Walker Boulevard
Augusta, Georgia 30912

706-721-7805 (Phone), 706-721-9799 (FAX)

Supplementary Table 1. The list of primers.

| Gene | Forward sequence | Reverse sequence |
|---|---|---|
| 18S | TCGAGGCCCTGTAATTGGAA | CCCTCCAATGGATCCTCGTT |
| ADRB3 | AGGCACAGGAATGCCACTCCAA | GCTTAGCCACAACGAACACTCG |
| CD206 | TCTTTGCCTTTCCCAGTCTCC | TGACACCCAGCGGAATTTC |
| DIO2 | GGTGGTCAACTTTGGTTCAGCC | AAGTCAGCCACCGAGGAGAACT |
| F4/80 | TCCTGCTGTGTCGTGCTGTTC | GCCGTCTGGTTGTCAGTCTTGTC |
| FGF21 | ATCAGGGAGGATGGAACAGTGG | AGCTCCATCTGGCTGTTGGCAA |
| GAPDH | AACTTTGGCATTGTGGAAGG | GGATGCAGGGATGATGTTCT |
| ICAM1 | AAACCAGACCCTGGAACTGCAC | GCCTGGCATTTCAGAGTCTGCT |
| IL6 | TACCACTTCACAAGTCGGAGGC | CTGCAAGTGCATCATCGTTGTTC |
| iNOS | ATGGACCAGTATAAGGCAAGC | GCTCTGGATGAGCCTATATTG |
| LEPRa | AATGACGCAGGGCTGTATGT | ATGGACTGTTGGGAAGTTGG |
| LEPRb | AATGACGCAGGGCTGTATGT | TCAGGCTCCAGAAGAAGAGG |
| LEPRe | TGAAGATGATGGAATGAAGTGG | AGTTGGCATCCTTATTTTGAGGT |
| NFIA | TGGACAGTCCTGGTGAAGAACC | CTTCAGTGTGCTTGGAGATGGC |
| NOX1 | CAGTTATTCATATCATTGCACACCTATTT | CAGAAGCGAGAGATCCATCCA |
| NOX2 | CAAGATGGAGGTGGGACAGT | GCTTATCACAGCCACAAGCA |
| NOX4 | TGTTGCATGTTTCAGGTGGT | AAAACCCTCGAGGCAAAGAT |
| Nrg4 | TCCTCCTCACTCTTACCATCGC | GTCTCTACCAGGCTGATCTCAC |
| PPARGC1A | GAATCAAGCCACTACAGACACCG | CATCCCTCTTGAGCCTTTCGTG |
| PRDM16 | ATCCACAGCACGGTGAAGCCAT | ACATCTGCCCACAGTCCTTGCA |
| TH | TGCACACAGTACATCCGTCATGC | GCAAATGTGCGGTCAGCCAACA |
| TNF-α | CCACCACGCTCTTCTGTCTAC | AGGGTCTGGGCCATAGAACT |
| UCP-1 | GCTTTGCCTCACTCAGGATTGG | CCAATGAACACTGCCACACCTC |
| VCAM1 | GCTATGAGGATGGAAGACTCTGG | ACTTGTGCAGCCACCTGAGATC |
| mt-16S rRNA | CCGCAAGGGAAAGATGAAAGAC | TCGTTTGGTTTCGGGGTTTC |
| mt-Nd1 | CTAGCAGAAACAAACCGGGC | CCGGCTGCGTATTCTACGTT |
| nu-B2m | TGGGAAGTCTAGGGAGGAGC | CATCCCATTCTGCACACCCT |
| nu-Rpph1 B | TTCGAGCCTTATAGTGGCGG | CAGTCTGCCCAAGCGTAGAG |

**Supplemental Figure 1.**

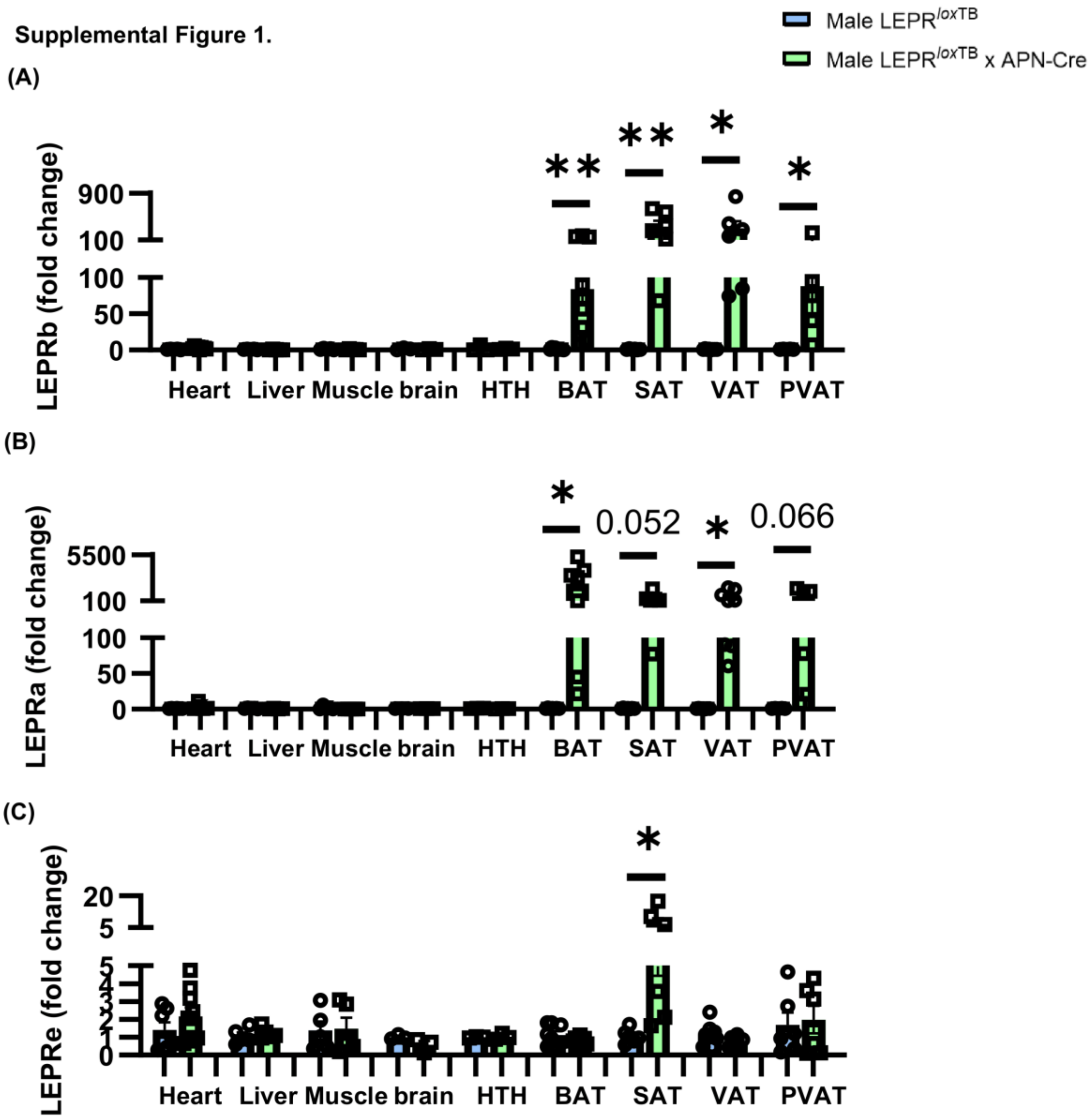


**Supplemental Figure 1. Restoration of adipocyte LEPR was specific to adipose tissues.** (A) Relative expressions assessed as each tissue of LEPR$^{loxTB}$ as a control by qPCR. Blue and green bars represent male LEPR$^{loxTB}$ and male LEPR$^{loxTB}$ x APN-Cre, respectively. Data are presented as mean ± SEM, n=3-9. *P<0.05, **P<0.01, ***P < 0.001.

Supplemental Figure 2.

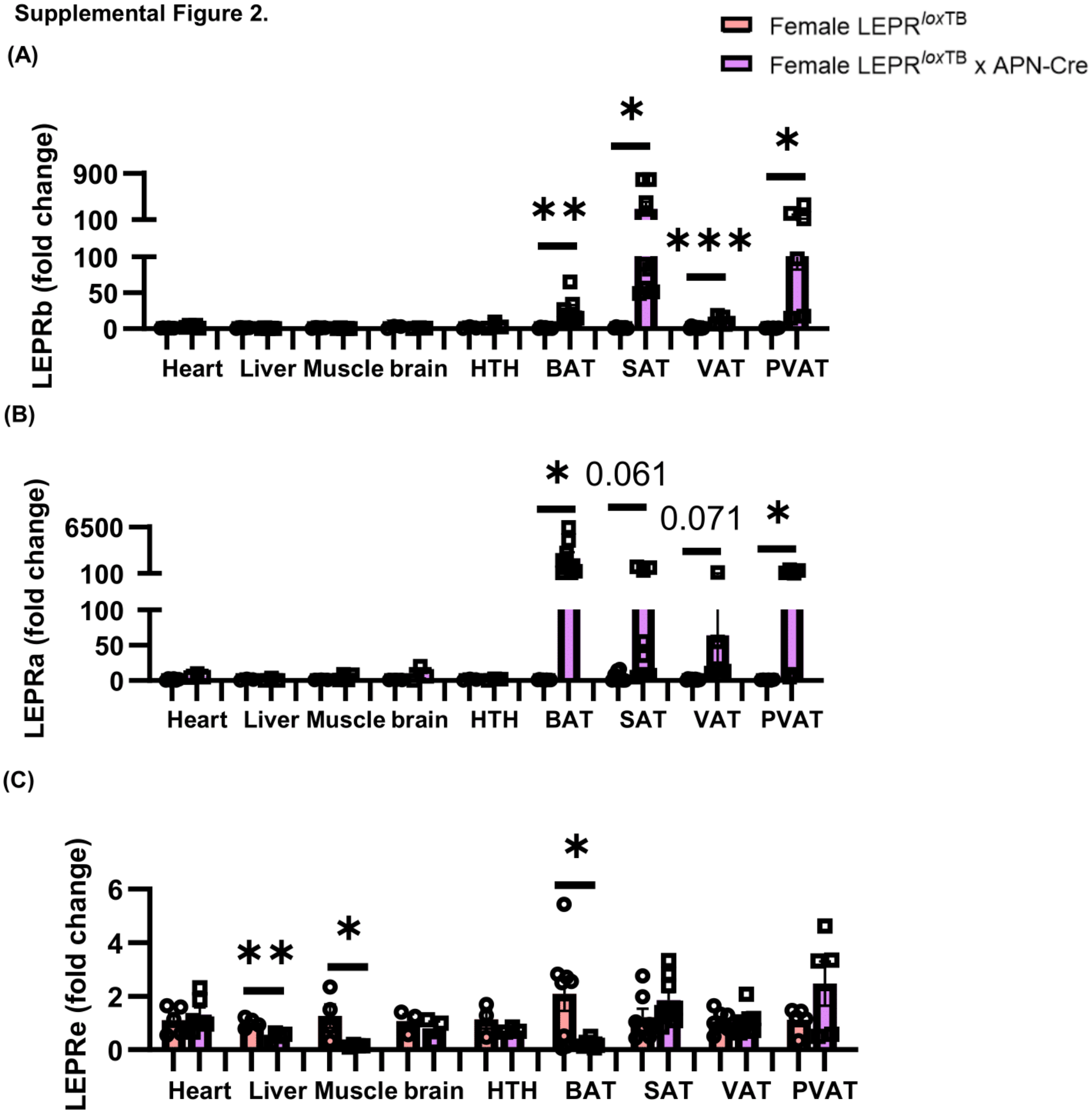


**Supplemental Figure 2. Restoration of adipocyte LEPR was specific to adipose tissues.** (A) Relative expressions assessed as each tissue of LEPR$^{loxTB}$ as a control by qPCR. Red and purple bars represent female LEPR$^{loxTB}$ and female LEPR$^{loxTB}$ x APN-Cre, respectively. Data are presented as mean ± SEM, n=3-9. *P < 0.05, **P < 0.01, ***P<0.001.

**Supplemental Figure 3.**

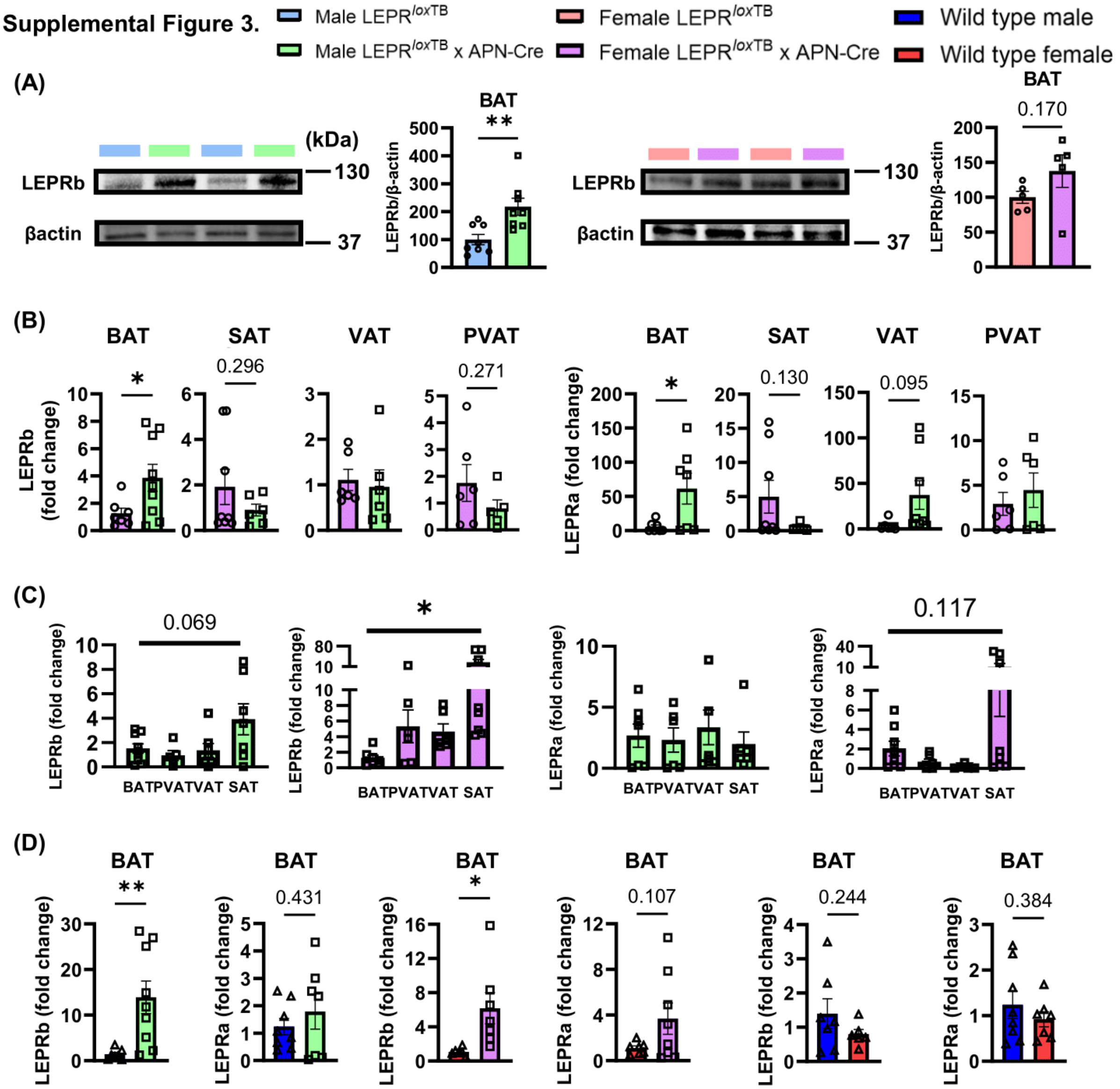


**Supplemental Figure 3. Restoration of adipocyte LEPR was more pronounced in BAT of male mice compared to females and that level was supraphysiological.** (A) Representative western blot and quantification of LEPR in male and female BAT. (B) Sex-based comparison of LEPR expression among adipose tissues, with female LEPR$^{loxTB}$ x APN-Cre mice used as reference. (C) LEPR expression levels among SAT, VAT, and PVAT in LEPR$^{loxTB}$ x APN-Cre mice normalized to BAT as the control in each sex. (D) LEPR expression levels among male and female LEPR$^{loxTB}$, LEPR$^{loxTB}$ x APN-Cre, and wild type in BAT. Blue, green, red, purple, dark blue, and dark red bars represent male LEPR$^{loxTB}$, male LEPR$^{loxTB}$ x APN-Cre, female LEPR$^{loxTB}$, female LEPR$^{loxTB}$ x APN-Cre, male wild type, and female wild type, respectively. Data are presented as mean ± SEM, n=5-9. *P < 0.05, **P < 0.01, ***P < 0.001.

**Supplemental Figure 4.**

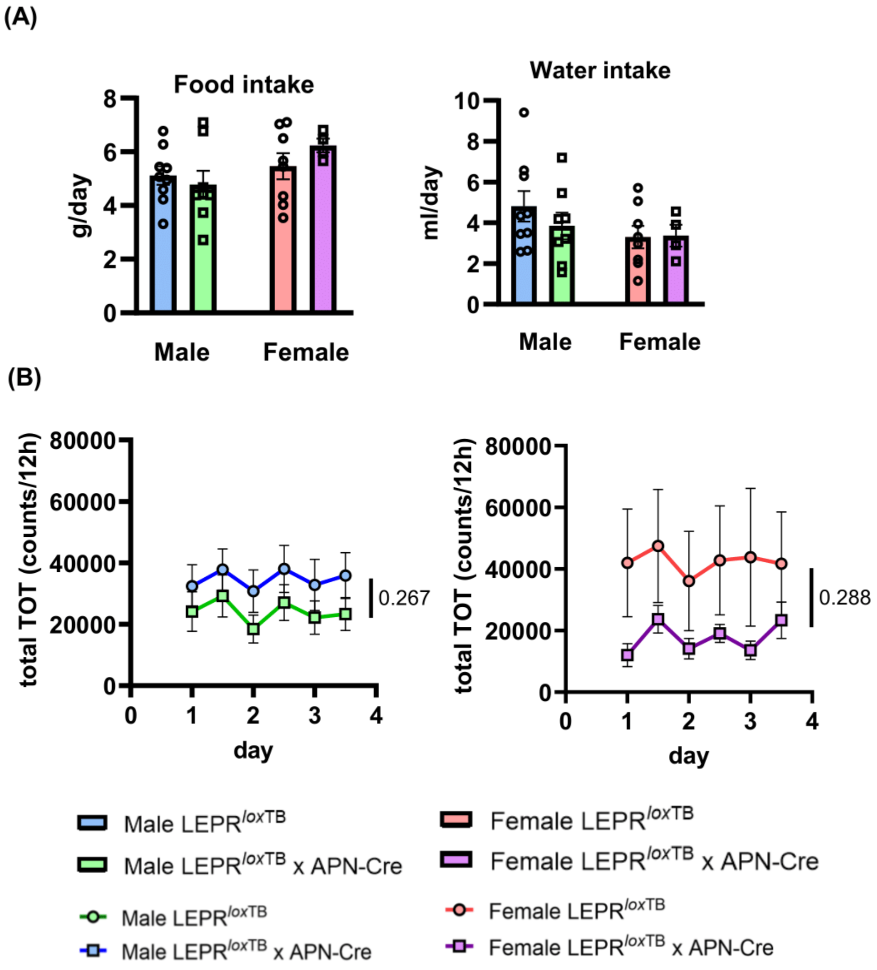


**Supplemental Figure 4. Restoration of adipocyte LEPR did not alter water and food consumption and activity.** (AB) Water and food consumption and activity among male and female LEPR$^{loxTB}$ and LEPR$^{loxTB}$ x APN-Cre. Blue, green, red, and purple bars represent male LEPR$^{loxTB}$, male LEPR$^{loxTB}$ x APN-Cre, female LEPR$^{loxTB}$, and female LEPR$^{loxTB}$ x APN-Cre, respectively. Data are presented as mean ± SEM, n=4-9.

**Supplemental Figure 5.**

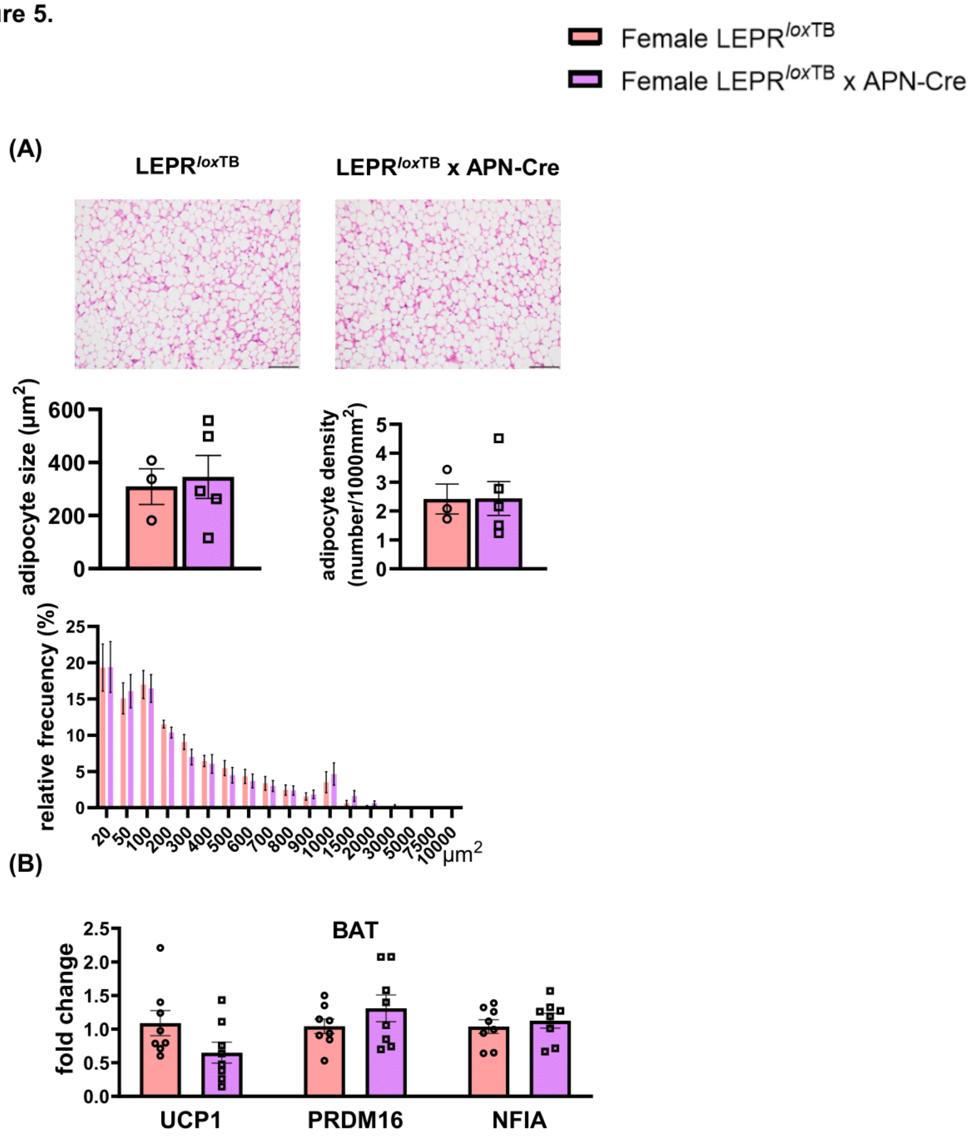


**Supplemental Figure 5. Restoration of adipocyte LEPR in females did not alter browning related genes nor size of adipocytes.** Representative section of BAT from female LEPR$^{loxTB}$ and LEPR$^{loxTB}$ x APN-Cre mice stained with H&E staining from which adipocyte size, density and frequency of distribution was measured. Scale bars indicate 50 μm (B) qPCR quantification of the browning markers, UCP1, PRDM16, and NFIA in female BAT. Red and purple bars represent female LEPR$^{loxTB}$ and female LEPR$^{loxTB}$ x APN-Cre, respectively. Data are presented as mean ± SEM, n=3-9. *P < 0.05.

**Supplemental Figure 6.**

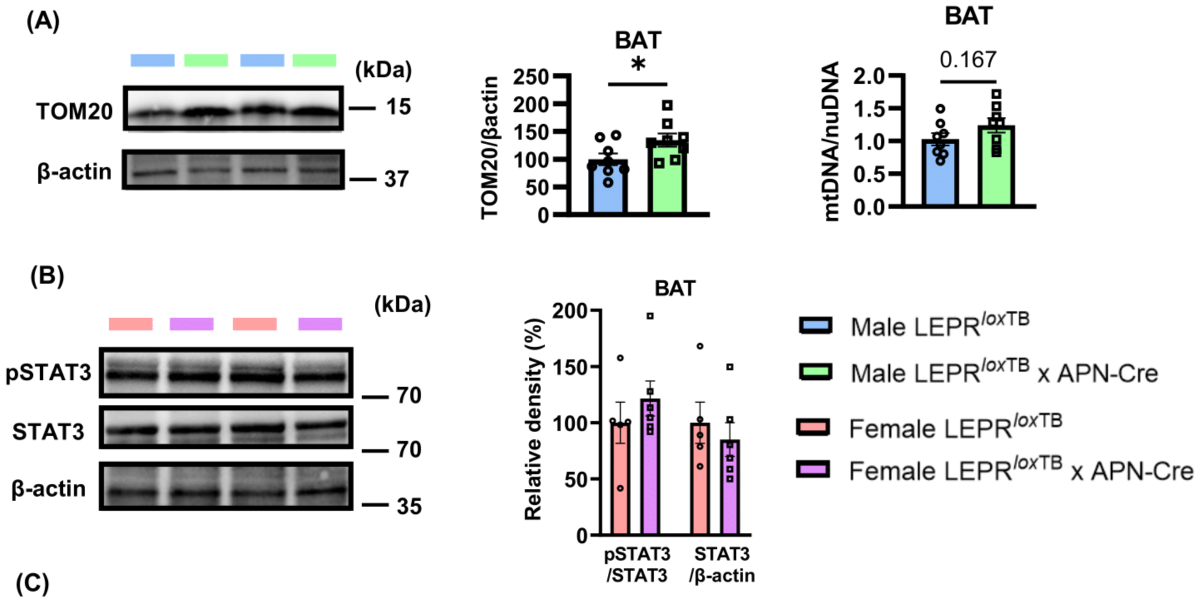


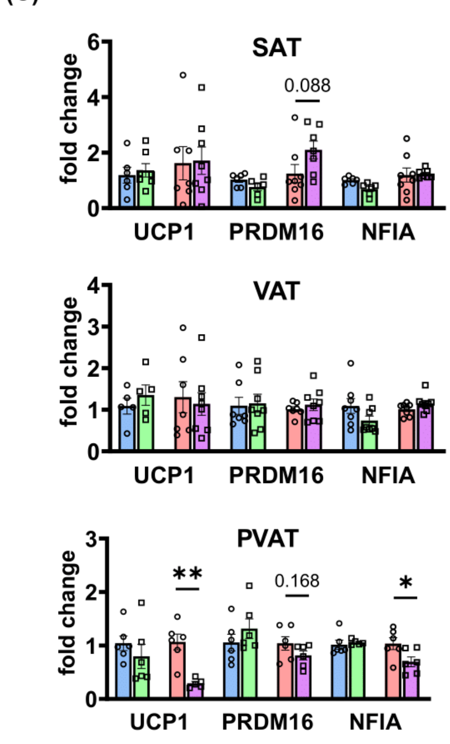


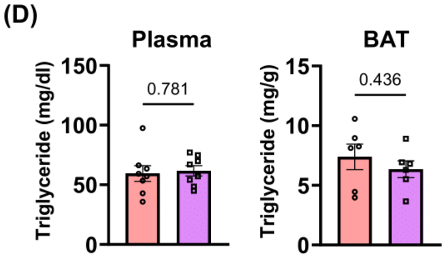


**Supplemental Figure 6. Restoration of adipocyte LEPR did not affect pSTAT3 phosphorylation in BAT of female mice, and it enhanced mitochondrial markers in BAT of male mice.** (A) Representative western blot and quantification of TOM20 and the mitochondrial-to-nuclear DNA ratio in male BAT. (B) Representative western blot and quantification of pSTAT3 and STAT3 in female BAT. (C) Comparison of the expression of browning-related genes in VAT, SAT, and PVAT in males and females, respectively. (D) Plasma and BAT triglyceride levels in females. Blue, green, red, and purple bars represent male LEPR$^{loxTB}$, male LEPR$^{loxTB}$ x APN-Cre, female LEPR$^{loxTB}$, and female LEPR$^{loxTB}$ x APN-Cre, respectively. Data are presented as mean ± SEM, n=5-8. *P < 0.05.

**Supplemental Figure 7.**

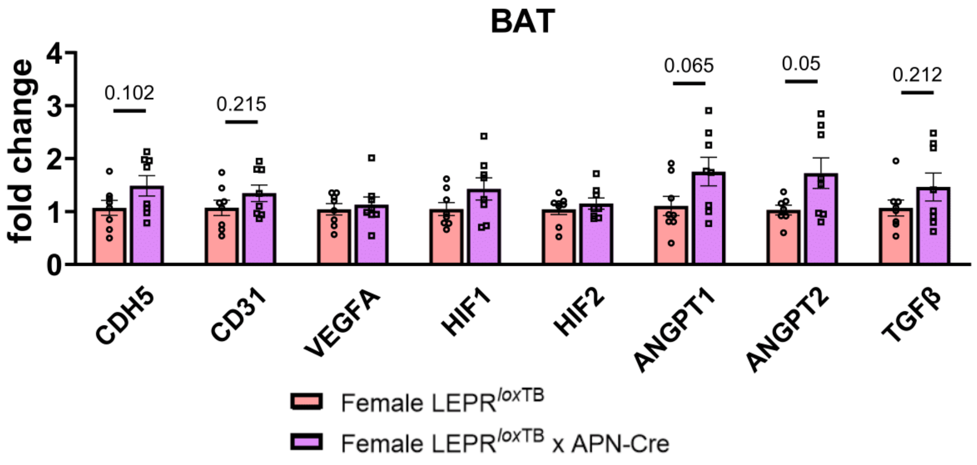


**Supplemental Figure 7. Restoration of adipocyte LEPR did not alter genes related to vascularization.** Red and purple bars represent female LEPR$^{loxTB}$ and female LEPR$^{loxTB}$ x APN-Cre, respectively. Data are presented as mean ± SEM, n=7-8. *P < 0.05, **P < 0.01, ***P < 0.001.

**Supplemental Figure 8.**

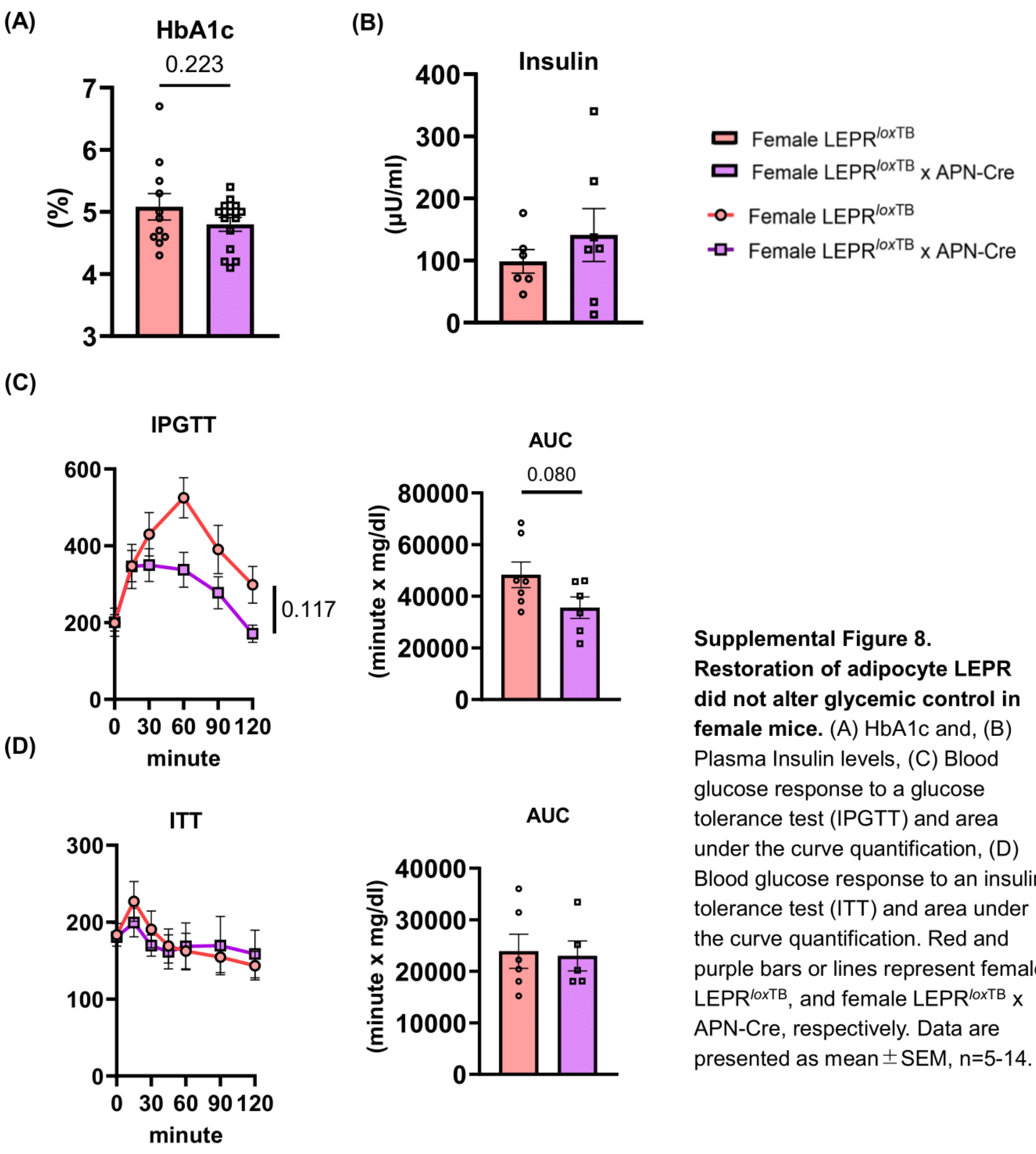


**Supplemental Figure 8. Restoration of adipocyte LEPR did not alter glycemic control in female mice.** (A) HbA1c and, (B) Plasma Insulin levels, (C) Blood glucose response to a glucose tolerance test (IPGTT) and area under the curve quantification, (D) Blood glucose response to an insulin tolerance test (ITT) and area under the curve quantification. Red and purple bars or lines represent female LEPR$^{loxTB}$, and female LEPR$^{loxTB}$ x APN-Cre, respectively. Data are presented as mean ± SEM, n=5-14.

**Supplemental Figure 9.**

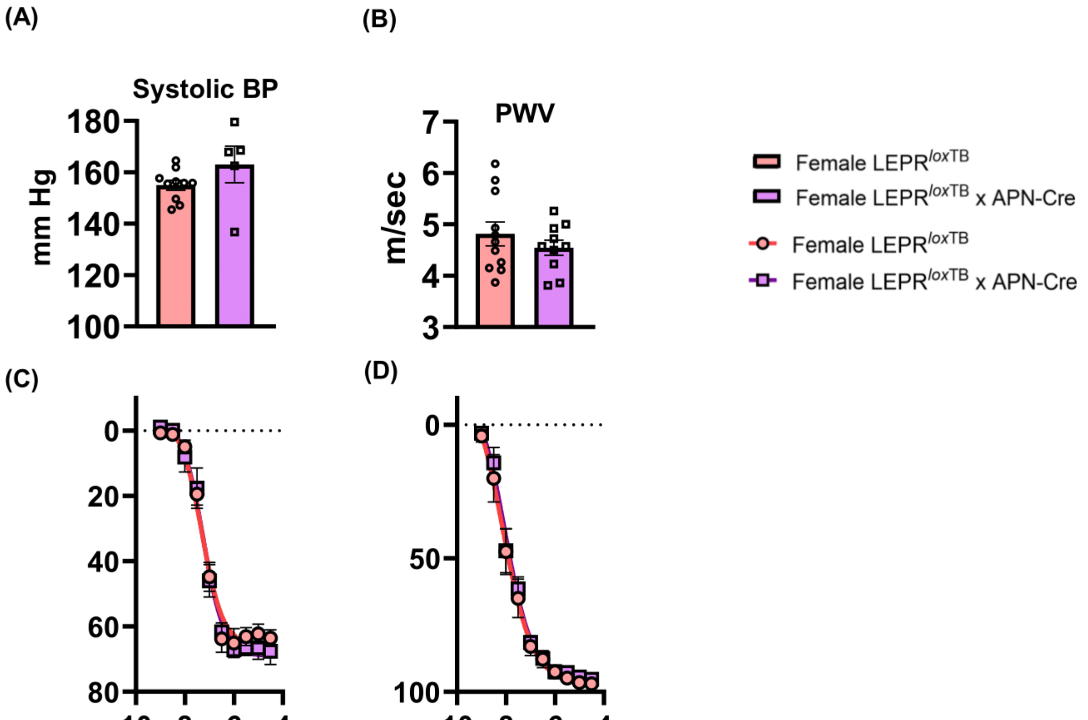


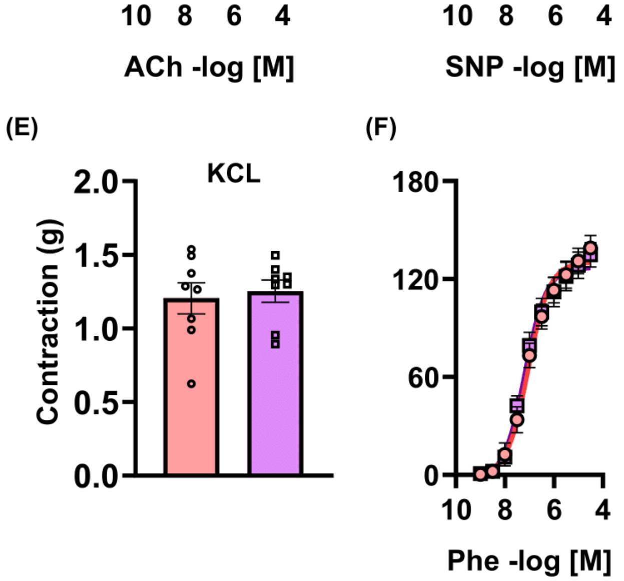


**Supplemental Figure 9. Restoration of adipocyte LEPR did not affect vascular function including blood pressure, arterial stiffness, and endothelial function in female mice.** (A) Systolic BP measured by tail-cuff method, (B) Pulse wave velocity (PWV), (C to F) Wire myograph analysis of aorta (C) endothelium-dependent, (D) endothelium independent relaxation, (EF) constriction to potassium chloride (80mM) and phenylephrine. Red and purple bars or lines represent female LEPR$^{loxTB}$ and female LEPR$^{loxTB}$ x APN-Cre, respectively. Data are presented as mean ± SEM, n=5-11.

**Supplemental Figure 10.**

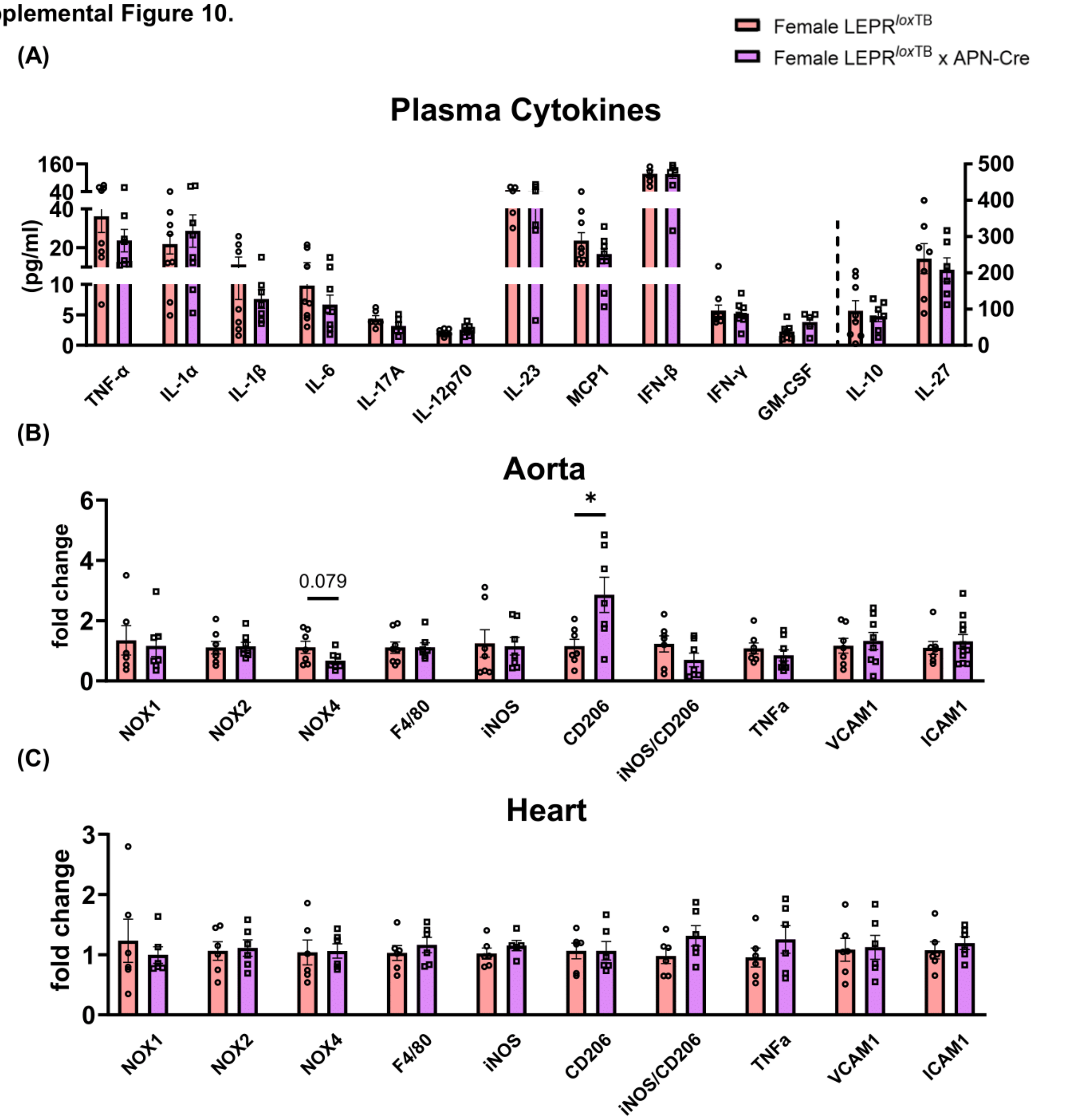


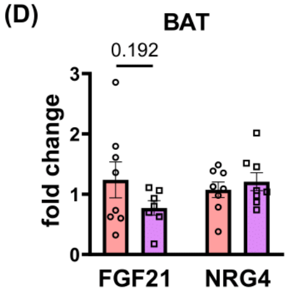


**Supplemental Figure 10. Restoration of adipocyte LEPR did not reduce local and systemic inflammation in female mice.** (A) Plasma cytokine, (B) Aorta and (C) Heart quantification of local inflammation via qPCR analysis. (D) qPCR quantification of FGF21 and NRG4 in BAT. Red and purple bars represent female LEPR$^{loxTB}$ and female LEPR$^{loxTB}$ x APN-Cre, respectively. Data are presented as mean ± SEM, n=5-11. *P < 0.05.
